# Interfacial water in the PRDX1–sulfiredoxin repair intermediate: an all-atom molecular dynamics study

**DOI:** 10.64898/2026.08.24.746838

**Authors:** Manal A. Nael, Laxman M. Alakonda, Eleonora Gianti, Khaled Elokely

## Abstract

Peroxiredoxins protect cells from oxidative damage, and sulfiredoxin (Srx) restores their activity by repairing the overoxidized catalytic cysteine; how the two proteins recognize one another is central to redox signaling and to oxidative-stress-associated disease. We characterize the human peroxiredoxin-1 (PRDX1)–Srx repair intermediate (PDB 2RII) and four related catalytic-cysteine states by all-atom molecular dynamics (three replicas of 200 ns per system; 3.0 µs total), using geometric and kinetic observables only, with across-replica statistics. The disulfide-linked complex is stable (backbone RMSD 2.5–3.2 Å) and the peroxidatic Cys52 is buried in the interface (buried surface area 71–76 Å²). The interface is extensive, with 202 consensus residue–residue contacts, and predominantly water-mediated: 20 reproducible, hydrogen-bond-validated water bridges centered on the PRDX1 165–170 region, against three persistent salt bridges. Removing the engineered tether and modeling Cys52 as the sulfinate on which Srx acts leaves the interface intact (189 consensus contacts) and the water network larger (36 bridges), new bridges linking the sulfinate to the Srx catalytic pocket. At the free Cys52, first-shell water responds to charge state as electrostatics predicts, and at matched water counts the thiolate shows no additional clustering, separating generic hydration from the specific interfacial organization. An equalized, permutation-controlled comparison of the Srx-bound and free intra- PRDX1 contact networks leaves them statistically indistinguishable: of 14,127 residue pairs only one exceeds both floors, and it does not reproduce across independent trajectory sets. A covalent celastrol–Cys173 adduct retains its thioether bond while the tethered ligand reorients widely, so covalent attachment fixes the anchor rather than the pose.

## 1. Introduction

Peroxiredoxins (Prxs) are abundant cysteine-dependent thiol peroxidases that reduce hydrogen peroxide, organic hydroperoxides, and peroxynitrite, and that act both as antioxidant enzymes and as regulators of peroxide-based signaling.[1, 2] In the typical 2-Cys Prx mechanism, the peroxidatic cysteine (C_P_) attacks the peroxide substrate and is oxidized to a sulfenic acid (CP- SOH). The resolving cysteine (C_R_) of the partner subunit then attacks the sulfenic sulfur, forming an intersubunit disulfide and releasing water. This disulfide is subsequently reduced by thioredoxin (Trx) to regenerate the active enzyme.[2, 3] Human PRDX1 (UniProt Q06830) is a prototypical 2- Cys Prx in which Cys52 is the peroxidatic cysteine and Cys173 the resolving cysteine.[4]

Under conditions of high oxidative flux, the sulfenic acid at C_P_ can be further oxidized to a cysteine-sulfinic acid (Cys-SO_2_H) before the resolving disulfide forms, inactivating the peroxidase.[4–6] This hyperoxidation underlies the "floodgate" model, in which transient inactivation of local peroxiredoxin allows hydrogen peroxide to accumulate and to participate in redox signalling.[3, 7] Sulfinic-acid formation was long thought irreversible until the discovery of sulfiredoxin (Srx; human SRXN1, UniProt Q9BYN0), an ATP-dependent enzyme that reduces the C_P_-sulfinic acid back to the thiol and so restores peroxidase activity.[8, 9] Srx-mediated repair is slow and mechanistically unusual, proceeding through phosphorylation of the sulfinate followed by a covalent Srx-Prx intermediate (Figure 1).[9, 10] This reversible regulation of PRDX1 activity is particularly relevant in cancer, where altered redox homeostasis creates a strong dependence on antioxidant systems that maintain reactive oxygen species (ROS) within a range compatible with tumor cell survival.[11]

**Figure 1.**
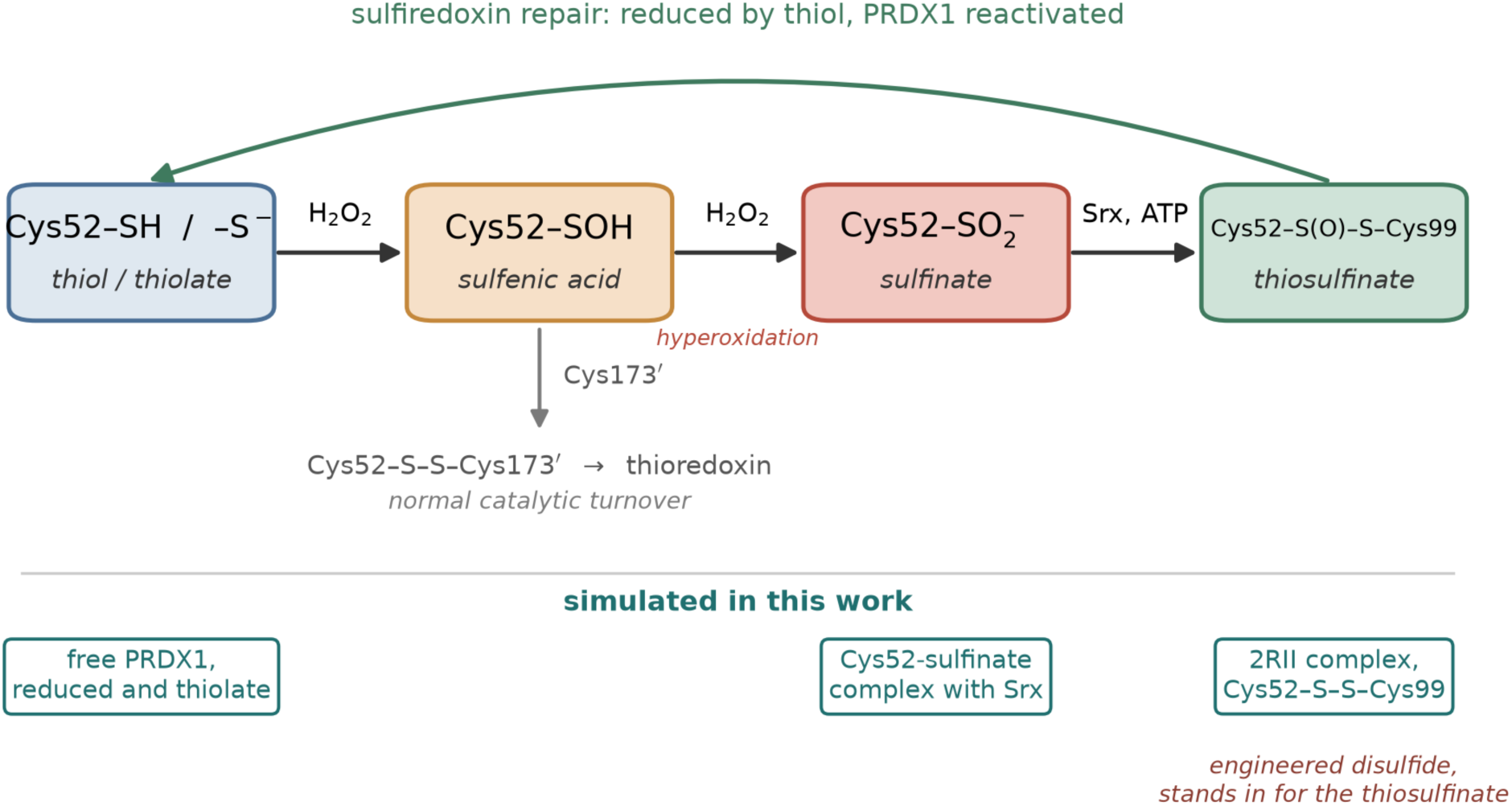
Oxidation states of the peroxidatic cysteine and the sulfiredoxin repair path. The peroxidatic Cys52 of PRDX1 cycles between the thiol/thiolate resting state and a sulfenic acid; in normal turnover the sulfenic acid condenses with the resolving Cys173 of the partner protomer to give an intersubunit disulfide that thioredoxin reduces. Under high peroxide flux the sulfenic acid is instead oxidized a second time to the sulfinate, which inactivates the peroxidase. Sulfiredoxin reverses this in an ATP-dependent reaction that proceeds through a sulfinic phosphoryl ester and then a covalent Srx–PRDX1 thiosulfinate, which is resolved by thiol to return Cys52 to the thiol state. Boxes at the foot of the scheme mark the states simulated here. Note that the crystallographic complex (PDB 2RII) carries an engineered Cys52–Cys99 disulfide in place of the labile thiosulfinate; the Cys52-sulfinate system simulated here corresponds instead to the state immediately preceding the covalent step.

One hallmark of cancer is disrupted redox homeostasis, resulting in elevated ROS driven by increased metabolic activity, mitochondrial dysfunction, oncogenic signaling, and rapid proliferation. To maintain ROS within a tolerable range, cancer cells rely heavily on antioxidant systems, including PRDX1, which detoxifies hydrogen peroxide (H₂O₂) through the thioredoxin-dependent catalytic cycle.[4] This dependence creates a potential therapeutic vulnerability. Tumors with elevated oxidative stress, including triple-negative breast cancer (TNBC),[11] may therefore exhibit increased reliance on PRDX1-mediated antioxidant defense. Understanding the molecular mechanisms that regulate PRDX1 activity and repair is essential for elucidating how redox homeostasis is maintained under oxidative stress.

Structural studies have established the molecular framework of Srx-mediated PRDX1 repair by trapping the PRDX1-Srx complex as a covalent intermediate (PDB 2RII).[12] Because the true sulfinic species is chemically labile, the complex was captured using a PRDX1 cysteine-substitution construct in which only the peroxidatic Cys52 remains native (Cys71 to Ser, Cys83 to Glu, Cys173 to Ser), linked to Srx through a mixed disulfide between PRDX1 Cys52 and the Srx catalytic Cys99. The 2.60 Å crystal structure shows an extensive "embrace" between the two proteins, but the solution dynamics of the interface, the disposition of the sequestered catalytic cysteine, and the contribution of ordered water to molecular recognition are not directly accessible from a single static model.

Beyond its established biochemical functions, the emerging therapeutic relevance of PRDX1, particularly in cancer, has positioned it as a potential redox-regulated therapeutic target, with several electrophilic small molecules reported to covalently modify its cysteine residues, highlighting the accessibility and chemical reactivity of redox-active cysteines as potential sites for therapeutic intervention.[13, 14] Celastrol, a pentacyclic triterpenoid quinone methide, forms a covalent thioether with PRDX1 Cys173 (PDB 7WET), providing a concrete structural model for covalent engagement of this protein.[14] How such a tethered adduct behaves in solution, in particular whether a persistent covalent bond implies a defined ligand pose, is directly relevant to covalent-inhibitor design.

Here we use all-atom molecular dynamics (MD) simulations (three independent 200 ns replicas per system; 3.0 µs in aggregate) with a staged-equilibration protocol and geometry- and kinetics-only observables to address five questions. (i) Is the trapped PRDX1-Srx complex stable over the simulated timescale, and how is the peroxidatic Cys52 disposed? (ii) Is the interfacial water organization specific recognition or generic hydration, and can it be separated from the charge-driven hydration of the free catalytic cysteine? (iii) Is the internal correlated-motion network of PRDX1 reproducible, and does it differ between the Srx-bound and free reduced states? (iv) Does the interface depend on the engineered disulfide, or does it survive when Cys52 is modeled as the sulfinate on which Srx actually acts? (v) Does the covalent celastrol-Cys173 adduct retain its bond, and how mobile is the tethered ligand? Throughout, the analysis focuses on observational quantities derived from molecular geometry and interaction kinetics, with uncertainty reported as the across-replica standard deviation (n = 3). We emphasize at the outset that the disulfide-linked complex (the PRDX1-Srx cysteine-trap intermediate) serves as a structurally accessible proxy for the native sulfinic-acid intermediate (hyperoxidized PRDX1 Cys52-SO₂H recognized by Srx), and that our findings should be interpreted within the context of the cysteine-trap construct used here.

## 2. Results

### 2.1. Stability of the complex and sequestration of the peroxidatic cysteine

The backbone RMSD (PRDX1-core aligned) reaches a plateau of 2.5-3.2 Å within ≤3.3 ns in all three replicas, and the individual chains remain folded (per-chain Cα root-mean-square-deviation, RMSD, last-half mean: PRDX1 A 1.6-2.6 Å, PRDX1 B 1.3-1.6 Å, Srx C 2.3-2.8 Å, Srx D 2.1-2.6 Å). The two mixed disulfides (PRDX1-Cys-S-S-Cys-Srx) stay formed (SG-SG 2.038 ± 0.002 Å across replicas; intact in every frame). They are, however, explicit bonds in the force-field topology and therefore cannot break during classical dynamics; the SG-SG distance is reported only to confirm that the bond term behaved normally and is not itself evidence of stability, which is instead read from the unconstrained observables above, namely the backbone RMSD and the per-chain folding (Fig. 2).

**Figure 2.**
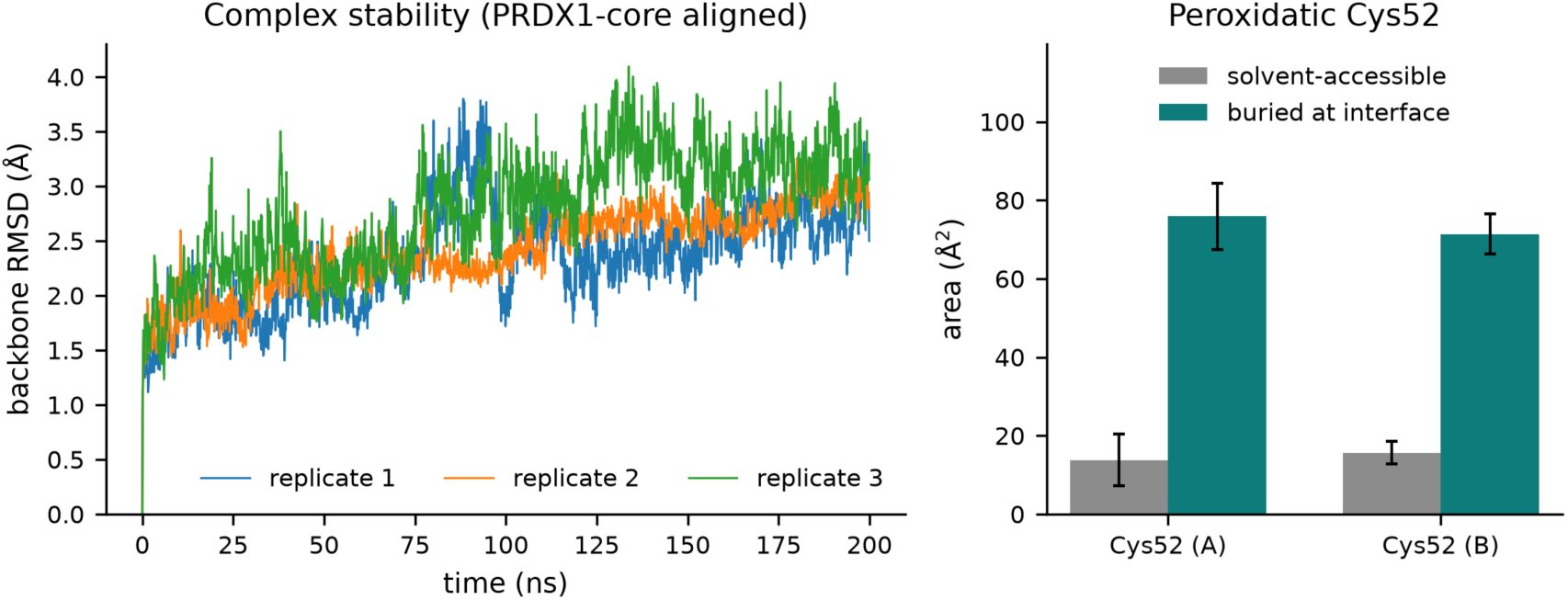
Stability of the trapped PRDX1–Srx complex. Backbone RMSD versus time for the PRDX1- Srx complex (PRDX1-core aligned) across three replicas, together with the solvent-accessible and buried surface area of the peroxidatic Cys52. RMSD is shown after periodic-image correction (anchored on a single disulfide-linked half).

The PRDX1-Srx interface is extensive, with 202 consensus residue-residue contacts (present in at least two of three replicas). The most persistent contacts (occupancy of 1.0 in all three replicas) involve PRDX1 Arg140, Asp146, Asp167, Thr166, Thr183, and Phe50 against Srx Ser132, His42, Asn43, Tyr128, and Gly130 (Table 1). The peroxidatic Cys52 is buried in this interface (solvent accessible surface area, SASA 13.8 and 15.6 Å² for protomers A and B; buried surface area 71-76 Å²; root-mean-square-fluctuation, RMSF 0.75-0.95 Å), disulfide-locked into the recognition surface. These observations are consistent with an extensive interface anchored by the PRDX1 C-terminal segment (Thr183, Pro148, Leu147) and the 140-167 region against a Srx surface centered on Ser132, with the catalytic cysteine sequestered (Fig. 3; Figure S1 and Figure S2).

**Figure 3.**
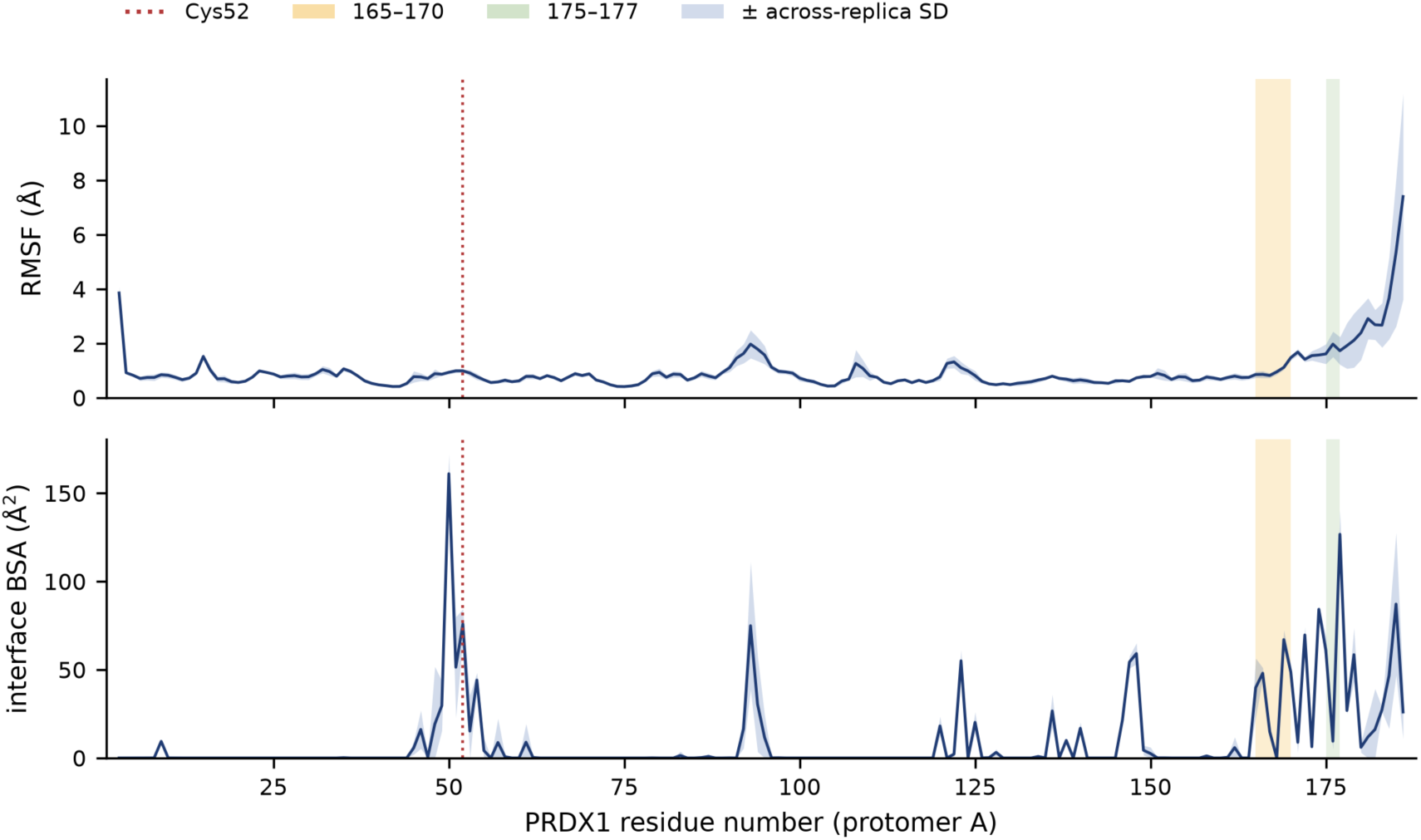
Per-residue flexibility and interface burial. Per-residue root-mean-square fluctuation (Cα, aligned per chain) and interface buried surface area (interface BSA) along the PRDX1 sequence; the 165- 170 recognition region is marked.

**Table 1.** Top persistent PRDX1-Srx interface contacts (occupancy across n = 3).

| PRDX1 | Srx | Occupancy (mean ± SD) |
| --- | --- | --- |
| Thr183 | His42 | 1.00 ± 0.00 |
| Asp167 | Ser132 | 1.00 ± 0.00 |
| Arg140 | Ser132 | 1.00 ± 0.00 |
| Asp146 | Ala131 | 1.00 ± 0.00 |
| Asp146 | Gly130 | 1.00 ± 0.00 |
| Thr183 | Asn43 | 1.00 ± 0.00 |
| Thr166 | Ser132 | 1.00 ± 0.00 |
| Pro148 | Leu129 | 1.00 ± 0.00 |
| Leu147 | Val127 | 1.00 ± 0.00 |
| Phe50 | Tyr128 | 1.00 ± 0.00 |
| Cys52 | Cys99 | 1.00 ± 0.00 (engineered disulfide — pipeline positive control; occupancy is 1.00 by construction) |

### 2.2. The PRDX1-Srx interface is water-mediated and specific

Our simulations reveal that the PRDX1/Srx repair interface is maintained through a dynamic network of hydrogen bonds, structured water molecules, and transient electrostatic interactions, highlighting the complementary roles of solvent-mediated and residue-level contacts in complex stabilization. Specifically, the interface contains 20 consensus persistent water bridges (hydrogen-bond-validated, identity-tracked, occupancy ≥0.5 in at least two of three replicas; per-replica 25, 21, and 26). These recur on both protomers and center on the 165-170 region and the Cys52- adjacent Phe50 (Table 2). By contrast, only two interface salt bridges are persistent and consensus (Lys185-Asp74, 0.71 ± 0.15; Glu123-Arg126, 0.75 ± 0.05). The interface is therefore predominantly water- and hydrogen-bond-mediated rather than salt-bridge-driven. These observations are consistent with specific, reproducible water-mediated recognition (named, persistent bridges that recur across protomers and replicas) at the 165-170 region and the Cys52- adjacent Phe50, as distinct from generic charge-driven hydration (Section 2.6; Fig. 5; Figure S3).

**Figure 4.**
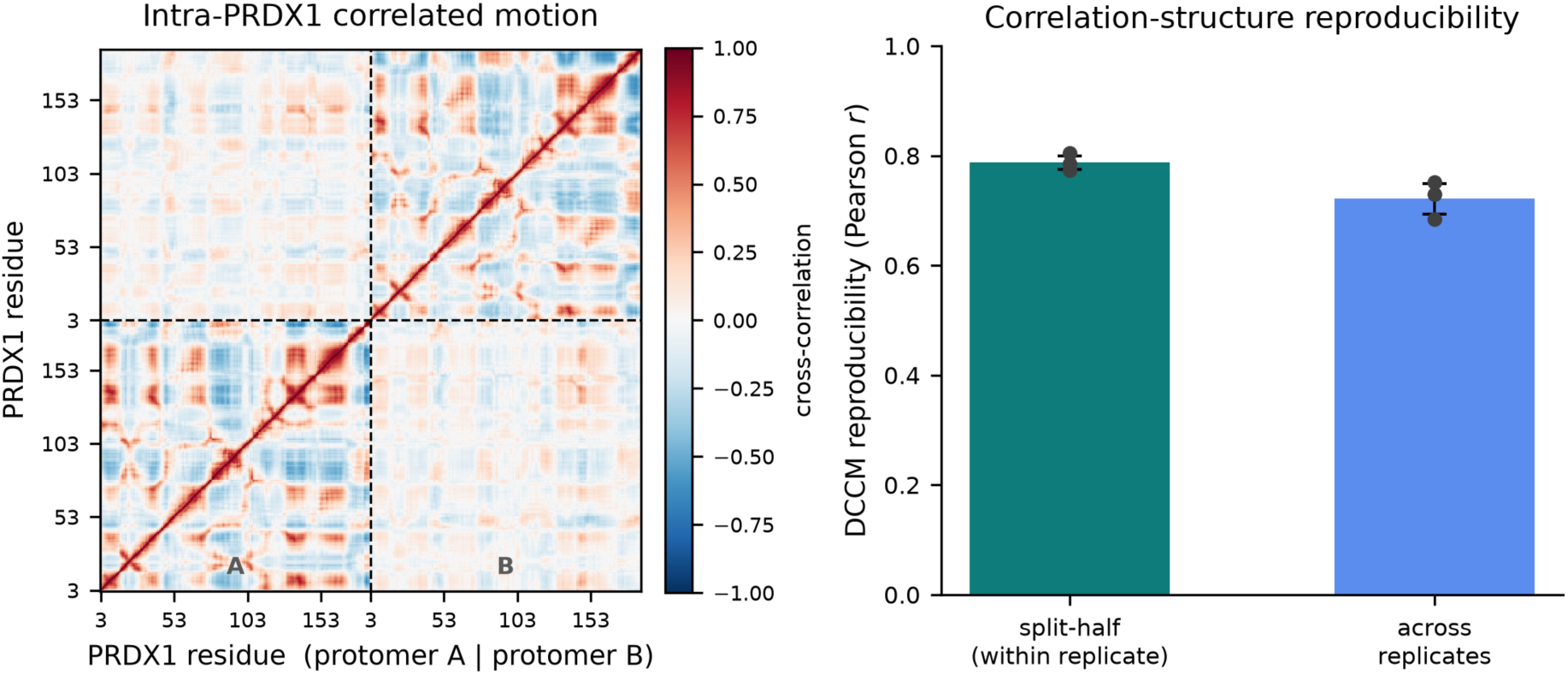
Internal correlated motion and network hubs. Across-replica mean intra-PRDX1 dynamic cross-correlation matrix and the betweenness-centrality hub ranking on the contact/DCCM graph. Hub identity is threshold-dependent (Section 2.9; Table S1).

**Figure 5.**
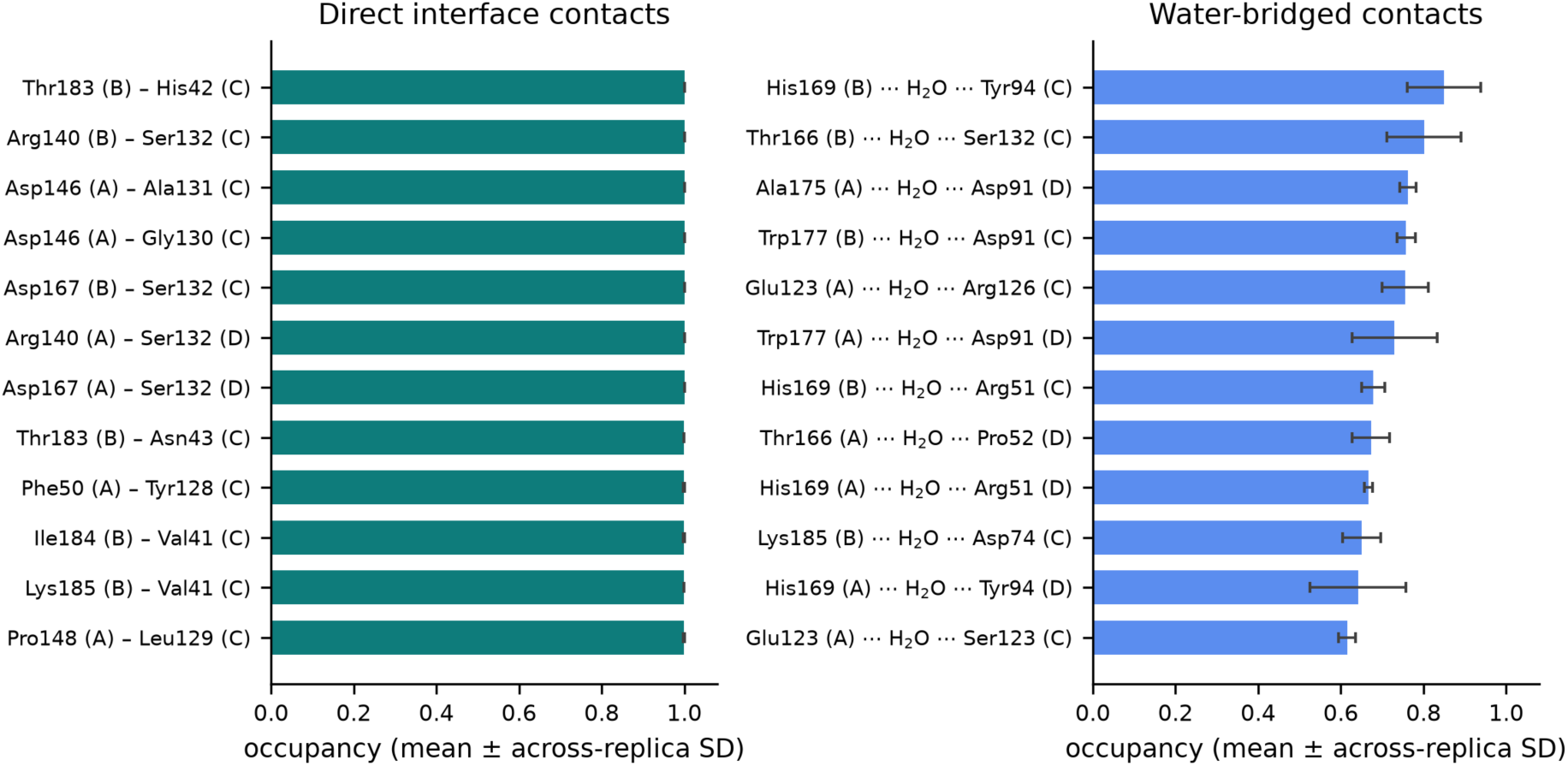
Interface contacts and persistent water bridges. The most persistent PRDX1-Srx residue- residue contacts and the consensus, hydrogen-bond-validated water bridges, with occupancy averaged across three replicas.

**Table 2.** Top consensus persistent interface water bridges (PRDX1 · H₂O · Srx)

| PRDX1 | Srx | Occupancy (mean ± SD) |
| --- | --- | --- |
| Thr166 | Ser132 | 0.90 ± 0.00 |
| His169 | Tyr94 | 0.89 ± 0.04 |
| Phe50 | Phe96 | 0.86 ± 0.02 |
| Phe50 | Asp80 | 0.85 ± 0.02 |
| His169 | Tyr94 (A) | 0.84 ± 0.14 |
| Phe50 | Gly97 | 0.81 ± 0.00 |
| Trp177 | Asp91 | 0.78 ± 0.11 |

*Each entry is a water molecule bridging the listed PRDX1 and Srx residues.*

### 2.3. The interface does not depend on the engineered tether

The complex characterized above is held together in part by a bond that does not exist in the native species: the crystallographic construct links PRDX1 Cys52 to Srx Cys99 through a mixed disulfide standing in for the labile thiosulfinate (Figure 1). This leaves a question the disulfide system cannot answer on its own — how much of the interface just described follows from that tether? We therefore built and simulated a system in which the tether is absent: Cys52 modeled as the sulfinate (–SO₂⁻), the chemical state on which Srx acts, with Srx Cys99 left as a free thiol. Construct, histidine states, box, protocol and analysis all match the disulfide complex, and the same three-replica, 200 ns treatment was applied.

The interface persists. The consensus contact count is 189, against 202 for the disulfide complex (per replica 191, 211 and 202), and the intra-PRDX1 contact network is essentially unchanged at 1546 versus 1556 consensus pairs. Backbone RMSD plateaus at 2.6–3.7 Å with a radius of gyration of 27.2–27.5 Å, against 2.5–3.2 Å and 26.8–27.5 Å for the disulfide complex; equilibration is slower, with per-replica plateau times of 4.0–5.8 ns against 1.2–3.3 ns.

The interfacial water network does not merely survive the change but expands. The sulfinate complex carries 36 consensus persistent water bridges (34, 33 and 37 per replica) against 20 for the disulfide complex, while consensus salt bridges fall from three to two. Compared protomer- agnostically, 11 of the 14 disulfide water-bridge residue pairs recur in the sulfinate (Jaccard index 0.46), among them every one of the most persistent: His169–Tyr94, Thr166–Ser132, Trp177– Asp91, Leu147–Tyr128 and Glu123–Arg126. Three pairs are absent (Asp146–Ala131, Lys185– Asp74, Thr166–Pro52) and ten are new (Table S5).

Four of the new bridges involve Cys52 itself. With the covalent bond removed, water occupies the space it vacated and bridges the sulfinate to the Srx catalytic pocket: Cys52···Cys99 at occupancy 0.66 ± 0.05, Cys52···His100 at 0.72 ± 0.05, and Cys52···Phe96 and Cys52···Gly98 at 0.51 ± 0.02 and 0.51 ± 0.01, each present in at least two of three replicas. Phe50 likewise gains bridges to Srx Asp80, Gly97 and Phe96. Over these trajectories the Cys52–Cys99 connection is therefore not lost when the covalent bond is taken away; it is re-formed through water.

Two limits on this reading. Clustering the pooled post-equilibration frames on the interface contact fingerprint gives a best silhouette of 0.127 at k = 2, so the ensemble does not separate into well-defined conformational substates and the cluster medoids are representative frames rather than distinct states. And the sulfinate group carries GAFF2 parameters for S(=O)O⁻ on an ff14SB backbone (Section 5.2), the same disclosure class as the celastrol junction; the system supports geometric and kinetic observables only.

### 2.4. Reproducible internal correlated motion with threshold-dependent hubs

The intra-PRDX1 dynamic cross-correlation matrix (DCCM) is reproducible: split-half 0.774- 0.804 within replica and across-replica 0.684-0.751 (Pearson correlation on the upper triangle). Betweenness centrality on the contact/DCCM graph identifies a recurring set of hubs in the PRDX1 hydrophobic core (Ala6, Tyr38, Ser71, Tyr116, Ile132, Ile133, and Leu139; top-15 in all replicas, pooled across the homodimer). However, the cutoff-sensitivity sweep (Section 2.9) shows that the identity of this hub set is threshold-dependent (Jaccard index[15] versus reference 0 to 1, median 0.125). The hub residues are therefore reported as a soft descriptor of the fold core rather than a fixed set; the converged, threshold-independent quantity is the DCCM reproducibility itself (Fig. 4).

### 2.5. Active-site pocket geometry

The active-site pocket nearest Cys52 is present in 100 of 100 sampled frames per replica, with volume 835 ± 184 Å³, 100 ± 20 alpha-spheres, apolar proportion 0.38 ± 0.03, hydrophobicity score 21.1 ± 3.3, polarity score 11.4 ± 1.9, and mean alpha-sphere radius 3.9 Å (geometric descriptors only; no druggability score is reported). The per-frame volume standard deviation is large, reflecting pocket breathing; the across-replica mean is reported.

### 2.6. Catalytic-cysteine hydration tracks charge state

At the free Cys52 SG, the thiolate draws roughly three times more first-shell water than the reduced thiol (1.37 to 4.41 within 3.5 Å), reorients the first-shell water dipoles toward the sulfur (orientation cosine +0.05 to -0.60), and draws in roughly three times more Na⁺ (0.11 to 0.38 within 8 Å) (Table 3). The thiolate system exhibits a marginal increase in the largest-cluster fraction (f_LC_ = 0.90 versus 0.87), but this difference does not persist when the two states are compared at equivalent first-shell water counts (Δf_LC_ = −0.11). The apparent increase therefore tracks the number of waters present rather than a change in the hydrogen-bonded network.

**Table 3.**
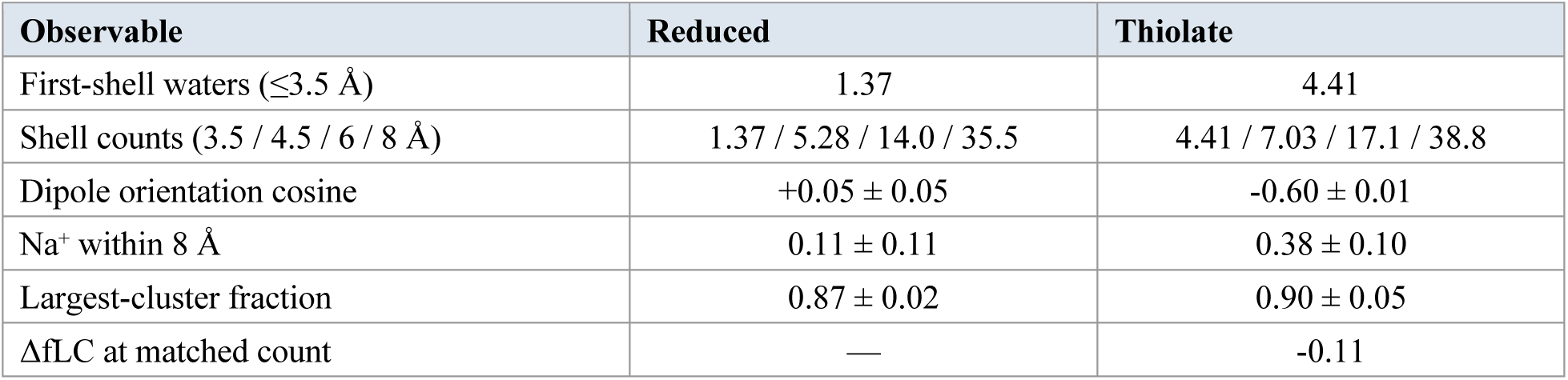
Cys52 hydration (reduced -SH versus thiolate -S⁻; across-replica mean ± SD).

These observations are consistent with an electrostatic hydration response (more water, dipoles oriented from hydrogen toward S⁻, more counterions) with no count-independent aggregation. Read together with Section 2.2, the two water populations behave differently: the water around the free catalytic cysteine responds to charge state as electrostatics would predict, whereas the waters at the Srx interface occupy specific positions and persist as identifiable bridges (Fig. 7; Figure S5).

**Figure 6.**
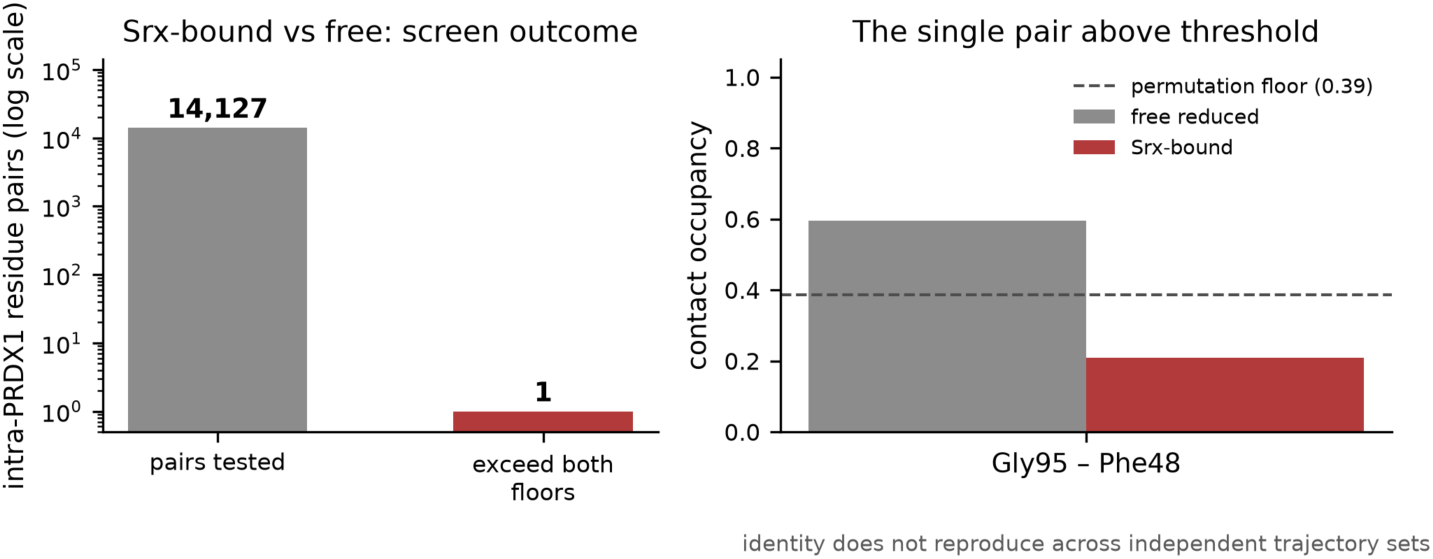
The intra-PRDX1 contact network is conserved between states. (Left) Of 14,127 intra- PRDX1 residue pairs screened in the Srx-bound versus free reduced comparison, one exceeds both the permutation null and the within-condition replicate floor. (Right) That pair (Gly95-Phe48) lies exactly at its permutation floor, and its identity does not reproduce across independent trajectory sets.

**Figure 7.**
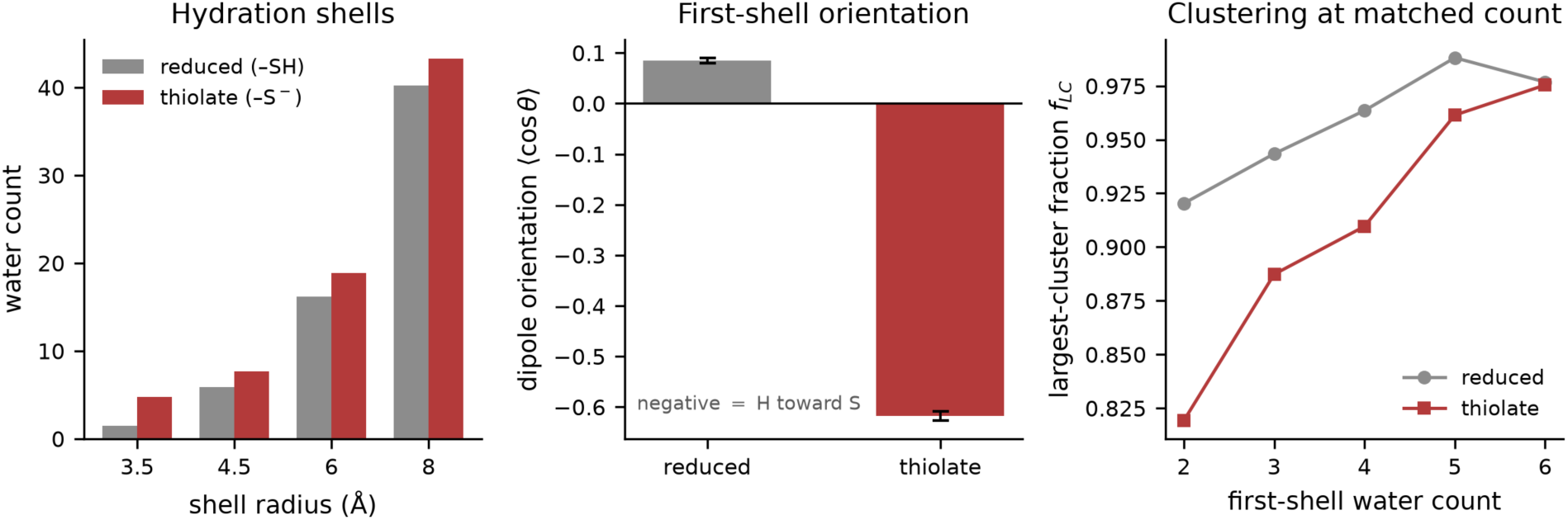
Charge-state dependence of catalytic-cysteine hydration. First-shell water counts, water- dipole orientation, and counterion occupancy at the free Cys52 SG for the reduced thiol and the thiolate, with a matched-water-count control.

### 2.7. A single inter-protomer contact distinguishes free and Srx-bound PRDX1

After equalizing the number of frames (1500 per state and replica), we compared 14,127 intra- PRDX1 residue pairs. Applying both a label-shuffle permutation test to assess statistical significance and a replicate-consistency threshold to exclude differences within the range of replica-to-replica variability identified only a single robust residue-pair difference (a second pair, Ile112-Phe43, differs by only Δ 0.03 and is near-saturated in both states). The flagged pair (Gly95- Phe48) shows an occupancy difference of 0.386, which coincides with its own permutation floor of 0.386, so it sits exactly at the detection threshold rather than clearly above it, and its occupancy is reduced rather than abolished (0.595 in the free reduced dimer, 0.209 in the Srx-bound complex; Table 4). Critically, the identity of that single above-threshold pair is not reproducible: an independent set of trajectories of the same systems, analyzed identically, returned a different single pair. What reproduces is the outcome of the screen, not its content. We therefore report the intra- PRDX1 contact network as statistically indistinguishable between the Srx-bound and free reduced states under this test, and assign no significance to any individual contact (Fig. 6; Figure S4). No causal or allosteric mechanism is inferred.

**Table 4.**
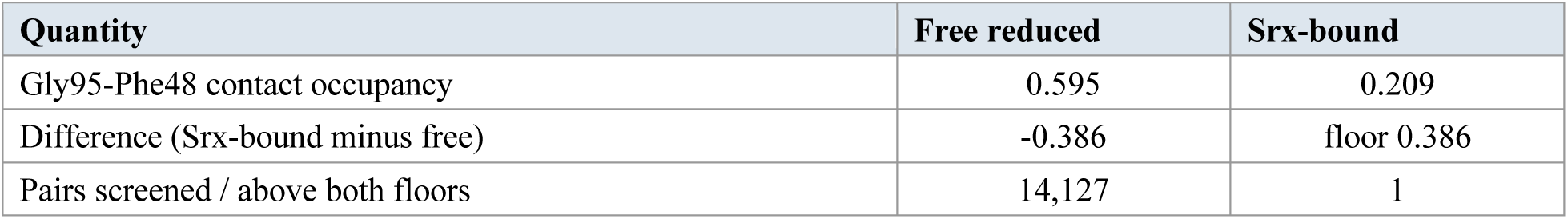
Outcome of the Srx-bound versus free reduced intra-PRDX1 contact screen.

### 2.8. The covalent celastrol adduct retains its bond but not its pose

The Cys173 SG-celastrol C6 thioether stays formed (1.821-1.825 ± 0.036 Å; intact in all replicas; maximum 1.97 Å) over 180 ns of production in three replicas. The ligand center of mass stays 4.5-5.0 Å from its anchoring SG (small SD), while the celastrol heavy-atom RMSD (protein-frame) differs systematically between the two protomers: 1.1-1.8 Å (maximum 4.8 Å) for the ligand on protomer A and 3.7-9.0 Å (maximum 13.6 Å) for the ligand on protomer B, reproducibly in all three replicas. These observations are consistent with the covalent bond fixing the attachment point but not the ligand pose, with the tethered ligand reorienting widely around the junction. The junction parameters are an approximate proxy, and this is a stability and mobility read rather than an energetic one (Fig. 8; Figure S6).

**Figure 8.**
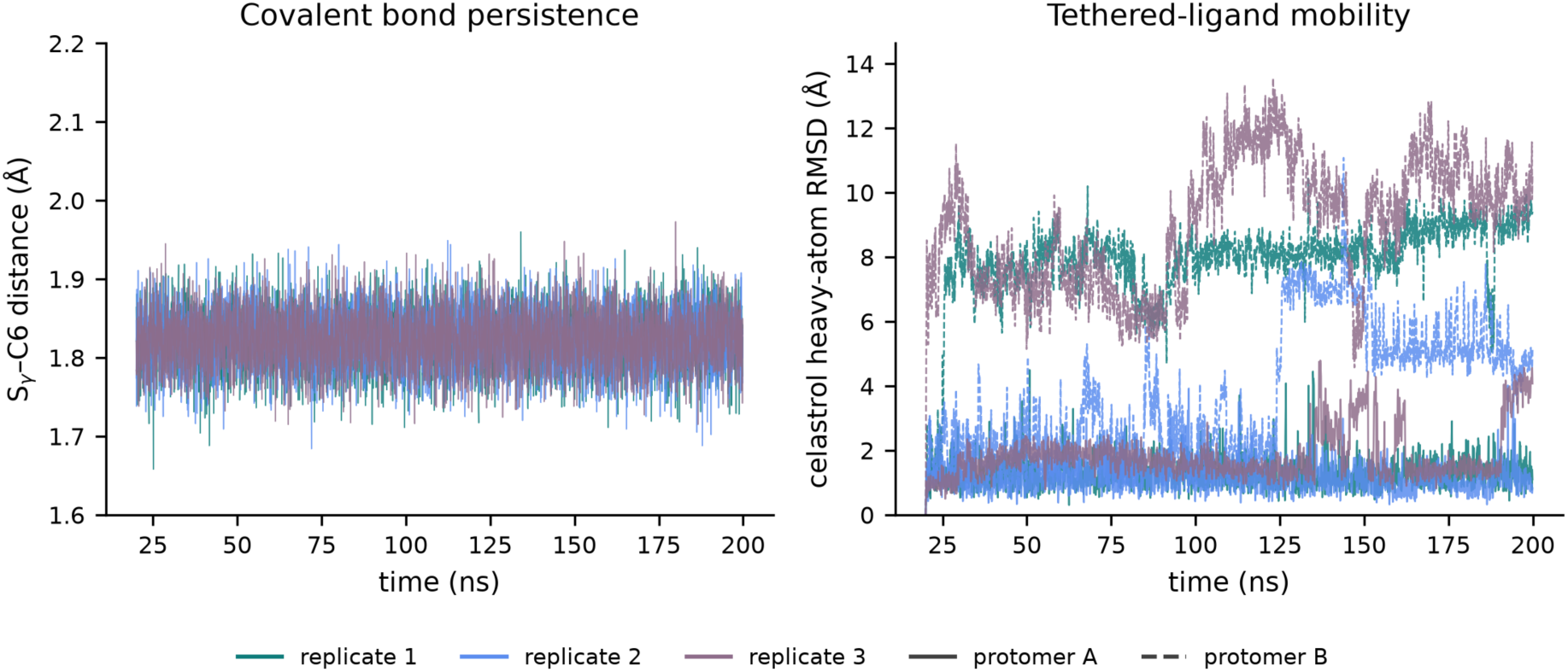
Behavior of the covalent celastrol adduct. Persistence of the Cys173 SG-celastrol C6 thioether bond and celastrol heavy-atom RMSD (protein-frame) across three replicas of the 7WET-derived system.

### 2.9. Convergence and threshold robustness

Over a 27-point grid (contact distance 0.40, 0.45, 0.50 nm; occupancy 0.4, 0.5, 0.6; |DCCM| 0.4, 0.5, 0.6), the interface consensus count ranges 83-214 (it scales with the threshold, as any count does; the occupancy-1.0 contacts of Table 1 are present throughout), and the betweenness hub set has a Jaccard index of 0 to 1 versus the reference (median 0.125) (Table S1). Per-replica backbone RMSD and radius of gyration plateau by ≤3.3 ns (a 20 ns discard is therefore conservative; per-replica equilibration times, last-half and final RMSD, per-chain RMSD, radius of gyration, and disulfide distances are tabulated in Table S2). The slow global observables are weakly sampled (effective sample size N_eff = 3.6-7.3 for backbone RMSD and 3.1-26.8 for radius of gyration; block standard error 0.17-0.22 Å and 0.04-0.17 Å, respectively), which is why across-replica standard deviation (n = 3) is the reported error throughout rather than within-trajectory statistics; the contact and interface observables sample faster than the global modes. Full per-system run parameters (versions, box and ion composition, histidine tautomers, disulfides, seeds, restraint schedule, and energies) are given in Table S3. The threshold-independent results (stability, interface contacts and Cys52 burial, the named water bridges, the catalytic-cysteine electrostatics, and the celastrol bond persistence) are the robust ones; the betweenness hub identity is threshold-sensitive (Table S1).

## 3. Discussion

The simulations give a coherent, geometry-level picture of the trapped PRDX1-Srx repair intermediate. The species studied is the cysteine-trap construct used to capture the complex, in which only the peroxidatic Cys52 is native and position 173 is a serine, and in which the genuine Cys-sulfinic intermediate that Srx acts on is represented by a tractable disulfide-linked proxy; the SG-SG distance is accordingly a force-field input rather than a measured observable. Within this construct, the complex is stable over the sampled timescale, the chains remain folded, the mixed disulfides remain near 2.0 Å, and the peroxidatic Cys52 is sequestered within an extensive recognition surface anchored by the PRDX1 C-terminal segment and the 140-167 region against a Srx surface centered on Ser132.

The central observation is that this recognition surface is predominantly water-mediated and specific. Twenty-three reproducible, hydrogen-bond-validated water bridges, named and recurring across protomers and replicas and centered on the 165-170 region and the Cys52-adjacent Phe50, organize the interface, whereas only two persistent salt bridges are present. Placing this beside the behavior of the free catalytic cysteine sharpens the interpretation: fixed-charge molecular dynamics captures the electrostatic and structural consequence of the cysteine protonation state, and on that footing the hydration of the free Cys52 is a straightforward electrostatic response, with the thiolate drawing more water, orienting water dipoles toward the sulfur, and recruiting counterions, but showing no aggregation beyond what its higher water count explains. The interfacial waters are thus specific recognition elements rather than generic charge-driven hydration, a distinction that is often blurred when water is discussed only qualitatively.

Comparing the Srx-bound and free reduced states, the intra-PRDX1 correlated-motion structure is reproducible and, under an equalized and floored comparison, largely conserved. Of 14,127 intra-PRDX1 residue pairs only one exceeds both floors, it lies exactly at its detection threshold, and its identity changes between independent trajectory sets analyzed identically. We therefore read the internal contact network as statistically indistinguishable between the two states rather than as carrying a specific state-dependent contact, and assign no significance to individual pairs. This reading is in any case bounded because the Srx-free reference is PRDX1 simulated from the decomplexed bound coordinates and oxidation state is therefore inseparable from Srx binding in this comparison.

Finally, the covalent celastrol-Cys173 adduct retains its thioether bond throughout while the tethered ligand reorients widely, with the heavy-atom RMSD reaching roughly 14 Å on one protomer. The junction here is described by approximate proxy parameters and is read for bond persistence and ligand mobility rather than for any energetic quantity, but on that read the message is clear: bond persistence does not imply a fixed ligand pose. For covalent-ligand design against PRDX1 and similar targets, this is a useful reminder that the warhead anchor and the reversible recognition of the rest of the ligand are separable, and that a stable covalent attachment can coexist with substantial conformational freedom of the appended scaffold.

Throughout, the analysis is deliberately observational and grounded in geometry and kinetics, enabling a direct characterization of structural features and dynamic behavior. The approach reports physically interpretable geometric pocket descriptors that capture key structural determinants without requiring free-energy, binding-affinity, kinetic-rate, or perturbation metrics. Because three replicas of 200 ns sample the slow global modes only modestly (effective sample sizes of a few to about ten for backbone RMSD and radius of gyration), across-replica standard deviation (n = 3) is used as the error unit, and the quantities emphasized here are the threshold-independent ones, namely the stability of the complex, the interface contacts and Cys52 burial, the named water bridges, the charge-state dependence of catalytic-cysteine hydration, and the persistence of the celastrol bond. These observations are specific to the simulated construct and to the crystallographic 2RII dimer[12] at 2.60 Å resolution from which they were initiated, and they are intended as a structural and dynamic reading of the repair intermediate rather than as energetic or mechanistic claims.

## 4. Conclusion

Our molecular dynamics simulations provide an atomistic view of the PRDX1–Srx recognition interface, revealing an extensive recognition surface in which 202 consensus residue–residue contacts are complemented by a reproducible network of structured, hydrogen-bond-validated interfacial water molecules, with three persistent salt bridges. At the free catalytic cysteine, the thiolate state increases local hydration and orients the first-shell water dipoles toward the sulfur, without producing clustering beyond what its higher water count accounts for; this charge-driven hydration is distinct from the specific, persistent water bridges that organize the Srx-bound interface. These findings refine the structural basis of sulfiredoxin-mediated repair of hyperoxidized PRDX1 and identify dynamic features of the interface that are not evident from static crystal structures. Given the central role of the PRDX1/Srx axis in maintaining redox homeostasis, particularly in triple-negative breast cancer (TNBC)[11] where PRDX1 and SRXN1 are frequently overexpressed to buffer elevated ROS levels, our results provide a mechanistic framework for understanding this interaction and establish a foundation for future efforts aimed at disrupting PRDX1–Srx recognition as a strategy to selectively impair antioxidant defenses in redox-dependent cancers.

## 5. Materials and Methods

### 5.1. Starting structures and constructs

Five systems were built (Table 5). The PRDX1-Srx complex was taken from PDB 2RII[12] (2.60 Å; PRDX1 chains A and B, Srx chains X and Y), with the two PRDX1 Cys52-Srx Cys99 mixed disulfides. Free reduced and free thiolate PRDX1 used chains A and B of 2RII with Cys52 as a reduced thiol or a deprotonated thiolate, respectively. The celastrol-Cys173 covalent adduct used PDB 7WET (PRDX1 chains A and B; celastrol, ligand AQR, thioether on Cys173).[14] The Cys52-sulfinate complex was derived from the same 2RII assembly with the engineered Cys52- Cys99 disulfides deleted, Cys52 converted to cysteine sulfinate (CSD, -CH₂-SO₂⁻) and Srx Cys99 left as a free thiol, so that the state immediately preceding the covalent step is represented rather than the trapped intermediate itself. The 2RII PRDX1 construct is a cysteine-substitution variant used to trap the repair intermediate: of the native cysteines only the peroxidatic Cys52 is present (Cys71 to Ser, Cys83 to Glu, Cys173 to Ser); all claims are therefore scoped to this construct, and the resolving Cys173 position is described as residue 173 rather than as a catalytic cysteine. Residues unresolved in 2RII (the PRDX1 N-terminus 1-2 and C-terminal tail 187+, and the Srx N-arm) are terminal, lie more than 20 Å from the interface, catalytic Cys52, and the 165-170 region, and were left unmodelled with default charged termini. Crystallographic waters and the phosphate ions in 2RII were removed and the systems re-solvated. PDB 4XCS (wild-type PRDX1 decamer; 2.10 Å)[16] was used only for active-site context illustration.

**Table 5.** Simulated systems and catalytic-cysteine states.

| System | PDB | Res. | Chains used | Catalytic-Cys state |
| --- | --- | --- | --- | --- |
| PRDX1-Srx complex | 2RII | 2.60 Å | A,B (PRDX1) + X,Y (Srx) | Cys52-Cys99 mixed disulfide (x2) |
| Free reduced PRDX1 | 2RII (A,B) | — | A,B | Cys52 reduced thiol |
| Free thiolate PRDX1 | 2RII (A,B) | — | A,B | Cys52 thiolate (CYM) |
| Celastrol-Cys173 adduct | 7WET | — | A,B | Cys173-celastrol thioether |
| Cys52-sulfinate complex | 2RII | 2.60 Å | A,B (PRDX1) + X,Y (Srx) | Cys52 sulfinate (CSD, x2); Srx Cys99 free thiol |

### 5.2. System preparation

Structures were prepared with the OpenMM tool PDBFixer[17] (missing heavy atoms rebuilt; non-standard residues replaced) and protonated at pH 7.4. Histidine tautomers were assigned by local geometry and cross-checked with PROPKA3[18] on the assembled complex: His10, His81, His84, and His169 (PRDX1) and Srx His34, His42, and His100 are neutral except His84, which was modeled as doubly protonated (HIP) in both PRDX1 protomers (PROPKA pKa 7.28 and 7.49; His84 forms a salt bridge with Asp47, Nε-Oδ 2.5-2.6 Å). His84 lies more than 14 Å from the interface and active site, so its protonation does not affect the reported observables. His169 (in the 165-170 region) and Srx His100 (at the interface) are confidently neutral (PROPKA pKa 6.0-6.4 and 5.4). The two PRDX1 Cys52-Srx Cys99 mixed disulfides were formed explicitly (exactly two cross-chain SG-SG bonds; zero in the free systems). The thiolate state was built by adding all hydrogens, then deleting the Cys52 Hγ and renaming CYS to CYM (deprotonated S⁻); the net charge is reflected in the counter-ion count. For the Cys52-sulfinate system the two sulfinate oxygens were placed on the Cys52 SG with S-O 1.47 Å, Cβ-S-O 104° and O-S-O 112°, the azimuth about the Cβ-S axis scanned over 720 orientations and chosen to direct the oxygens away from Srx (closest O to Srx 2.98 Å); deleting the disulfide leaves a 1.55 Å S···S contact, relieved by rotating the Srx Cys99 χ1 torsion alone, with all backbones untouched, to give S···S separations of 3.4-3.7 Å. Sulfinate charges were derived with AM1-BCC on a capped ACE-CSD-NME fragment (net -1). Because ff14SB carries no atom type for a three-coordinate sulfinate sulfur, the S(=O)O⁻ group was assigned GAFF2 types (sulfur s4, oxygens o) on an otherwise ff14SB backbone; this is the same disclosure class as the celastrol junction and, as there, the system is used for geometric and kinetic observables only, never for an energetic quantity. Celastrol (AQR in 7WET[14]) was parameterized with GAFF2 and AM1-BCC charges[19–21] (antechamber and parmchk2, AmberTools); the Cys173 SG-celastrol C6 thioether was formed in tleap with explicit, approximate proxy bond, angle, and dihedral terms for the junction, and ff14SB[22] was used for the protein. This proxy junction supports a covalent-adduct stability and mobility analysis only and is not used for any energetic quantity.

### 5.3. Solvation, force field, and integration

Systems were solvated in a cubic TIP3P[23] box with 1.2 nm padding and 0.15 M NaCl, neutralized to the correct per-state net charge (complex: 124 Na⁺, 120 Cl⁻, 43,993 waters; thiolate: two additional Na⁺ for the net charge). Protein and water used amber14-all.xml and amber14/tip3p.xml (OpenMM[17]) or ff14SB[22], GAFF2[19], and TIP3P[23] (S6 adduct; AmberTools and tleap)[24]. Particle-mesh Ewald[25] was used (1.0 nm real-space cutoff; Ewald tolerance 5 x 10⁻⁴), with HBonds constraints and rigid water and a 2 fs timestep. The temperature was 310.15 K (LangevinMiddle integrator; 1 ps⁻¹ friction) and the pressure 1 bar (Monte Carlo barostat; 25-step interval). Integrator and barostat random seeds were recorded in per-replica manifests. Simulations used OpenMM 8.5.1[17] and Python 3.11.[26]

### 5.4. Equilibration and production

Each replica followed a staged equilibration before the production clock: (1) minimization to convergence (L-BFGS; tolerance 10 kJ mol⁻¹ nm⁻¹); (2) restrained NVT heating 10 to 310.15 K over 0.3 ns with harmonic positional restraints (k = 1000 kJ mol⁻¹ nm⁻²) on all protein heavy atoms; (3) restrained NPT for 2 ns at 310.15 K and 1 bar; (4) stepwise restraint release over 2 ns (k = 1000, 500, 250, 100, 50, 10, 0 kJ mol⁻¹ nm⁻²); (5) free NPT for 5 ns (unrestrained; not analyzed); and (6) production for 200 ns as a single continuous trajectory (no velocity reset), with frames every 20 ps. Three independent replicas per system (different integrator and barostat seeds) were run for the complex, free reduced, free thiolate, and S6 adduct systems (200 ns x 3 each; 12 production trajectories; 2.4 µs aggregate). Disulfides were verified intact after equilibration and after production. For the S6 amber-prmtop system, minimization was run unrestrained first, then heavy-atom restraints were applied to the minimized coordinates.

### 5.5. Analysis

A shared module enforced three corrections across every analysis: (I) periodic-image imaging (molecules made whole and the complex re-imaged anchored on a single disulfide-linked PRDX1- Srx half, because the two halves are separate molecules that otherwise drift into different periodic images and inflate apparent RMSD). Prior to imaging, whole-complex backbone RMSD reached 46–83 Å (last-half mean 36–45 Å across replicas), whereas the chains remained internally folded and the disulfide linkages were maintained at ∼2.0 Å; after anchoring imaging on one half, the same quantity is 2.5–3.2 Å; (II) a production-equilibration discard (the first 20 ns of production removed, justified by per-replica backbone-RMSD and radius-of-gyration plateaus, so no observable references frame 0); and (III) a canonical residue map derived geometrically from the catalytic-cysteine landmark so that the topology renumbering is consistent in all labels. The imaged trajectories were then checked directly for minimum-image contact: for 400 evenly spaced frames per replica (every 500 ps of production), the shortest distance between any solute atom and any periodic image of the solute was computed over all 26 neighboring lattice translations. For the free reduced, free thiolate, celastrol-adduct and Cys52-sulfinate systems this distance stays at or above 22.8 Å in every frame examined, so no frame reaches the 10 Å real-space cutoff. For the disulfide-linked complex the closest approach is 7.5 Å, in replica 1, where 1.3% of frames fall below the cutoff (0.4% across the three replicas); replicas 2 and 3 stay above it throughout (Table S4).

Per-residue analyses comprised RMSF (Cα, aligned per chain), SASA and per-residue buried surface area (BSA = SASA(isolated chains) - SASA(complex); Shrake-Rupley[27]), DSSP secondary-structure propensity,[28] hydration (waters within 4.5 Å), and intra-protein contact degree. Residue-residue contacts used the closest heavy-atom distance below 0.45 nm (candidate pairs pre-filtered by mean-Cα proximity); persistent contacts had occupancy ≥0.5, and interface contacts were reported as consensus (present in at least two of three replicas; mean occupancy ± SD). Correlated motion used the dynamic cross-correlation matrix on the intra-PRDX1 Cα subgraph (per-chain-aligned displacement field), with split-half and across-replica reproducibility reported; betweenness centrality used a graph with edges of occupancy ≥0.5 and |DCCM| > 0.5, pooled across the homodimer. Interface water bridges were defined as a water whose oxygen is within 0.35 nm of a PRDX1 polar (N or O) atom and a Srx polar atom and that donates a geometric hydrogen bond (angle ≥120 degrees; H-acceptor ≤0.25 nm) to at least one side, tracked by residue- pair identity; persistent water bridges had occupancy ≥0.5 and consensus in at least two of three replicas. Salt bridges used a charged side-chain N-O distance ≤0.40 nm, deduplicated to residue pairs. Pockets were detected with fpocket[29] (at least 100 frames); the pocket nearest the catalytic Cys52 SG (within a 5 Å ceiling) was reported with geometric descriptors only (the druggability score excluded). The Srx-bound versus free comparison used intra-PRDX1 contact occupancy per protomer-agnostic residue pair, equalized to 1500 frames per state per replica; a pair was reported as differing only if |Δ occupancy| exceeded both a label-shuffle permutation null (95th percentile; 3000 permutations) and the within-condition replicate-difference floor (95th percentile) and was persistent (occupancy ≥0.5) in at least one state. Catalytic-cysteine water (reduced versus thiolate) was characterized at each Cys52 SG by shell water counts (3.5, 4.5, 6.0, 8.0 Å), first-shell water- dipole orientation (cosine of the water dipole versus the SG-to-O vector; negative indicates hydrogen toward sulfur), water clustering (largest-cluster fraction and O-O coordination, O-O < 0.35 nm), and counterion shells, with clustering compared between states within each first-shell water-count bin (matched-count control). The S6 adduct was analyzed by SG-C6 bond distance versus time, celastrol heavy-atom RMSD after protein-backbone alignment, and ligand center-of-mass excursion from its anchoring SG. Cutoff sensitivity was assessed by recomputing the interface consensus count and the betweenness hub set over contact distances of 0.40, 0.45, and 0.50 nm, occupancies of 0.4, 0.5, and 0.6, and |DCCM| of 0.4, 0.5, and 0.6. Convergence was assessed from per-replica backbone-RMSD and radius-of-gyration plateaus, block standard error, and effective sample size (integrated autocorrelation); across-replica standard deviation (n = 3) is the reported error for all means. No free-energy, binding-affinity, kinetic-rate, or perturbation quantities were computed at any stage of this work.

### 5.6. Software

Software comprised OpenMM 8.5.1,[17] AmberTools (antechamber, parmchk2, tleap),[24] PDBFixer,[17] PROPKA3,[18] MDTraj 1.11,[30] MDAnalysis 2.10,[31] NetworkX 3.6,[32] scikit-learn,[33] SciPy,[34] NumPy[35], and fpocket.[29] Per-replica run manifests (versions, force-field files, seeds, ion counts, histidine tautomers, and energies) accompany the deposited data.

## Data availability statement

The experimental starting structures used to build every system are publicly available in the Protein Data Bank under accession codes 2RII, 4XCS, and 7WET. All data supporting the reported findings are contained within the article and its Supporting Information. The analysis procedures are specified in full in the Materials and Methods and, in implementation-independent algorithmic form with every threshold and control stated, in the Supporting Information; derived data are available from the corresponding author on reasonable request.

## Author contributions

**M.A.N.:** Conceptualization; Methodology; Software; Validation; Formal analysis; Investigation; Data curation; Visualization; Writing – original draft; Writing – review and editing. **L.M.A.:** Methodology; Validation; Investigation; Formal analysis; Writing – review and editing. **E.G.:** Conceptualization; Validation; Funding acquisition; Writing – review and editing. **K.E.:** Conceptualization; Methodology; Software; Supervision; Project administration; Resources; Funding acquisition; Writing – review and editing. All authors have read and agreed to the submitted version of the manuscript.

## Acknowledgments

Artificial-intelligence tools were used to assist with literature searching and language editing. All scientific content, analyses, interpretations, and conclusions are the authors’ own.

## Funding Sources

This publication was made possible by an Institutional Development Award (IDeA) from the National Institute of General Medical Sciences of the National Institutes of Health under Grant # 2P20GM103432 (K.E.). The content is solely the responsibility of the authors and does not necessarily represent the official views of the National Institutes of Health.

This research was supported by The RF Research Foundation and the City University of New York (CUNY) Queens College startup funds (EG); the PSC-CUNY Award # GR-00017761 (EG) and the CUNY Faculty Fellowship Publication Program (FFPP) Office of Faculty Affairs, The City University of New York (EG); the University of Wyoming (UWYO) and School of Pharmacy (SOP) startup funds (K.E.).

Computational resources were provided by the Advanced Research Computing Center (ARCC) at the University of Wyoming.

## Supporting information

Supporting information is provided in a separate file (Supplementary_Information).

Supporting figures and tables use the prefix S and are cited at the relevant points in the main text. The Supporting Information contains: Figure S1, the 2RII four-chain assembly; Figure S2, the interface around the buried catalytic Cys52; Figure S3, the 165-170 water-recognition site; Figure S4, the Srx-bound versus free intra-PRDX1 contact screen contact; Figure S5, the catalytic-cysteine environment; Figure S6, the celastrol-Cys173 covalent adduct; Table S1, the cutoff- sensitivity grid (interface consensus count and betweenness-hub Jaccard over the 27-point threshold grid); Table S2, per-replica convergence diagnostics; Table S3, per-system run manifests (versions, box and ion composition, histidine tautomers, disulfides, seeds, restraint schedule, and energies); Table S4, the minimum-image separation measured over all 26 lattice translations; and Table S5, the consensus interface hydrogen-bond network compared between the disulfide-trapped and Cys52-sulfinate states.

