## Supplementary material for "Interfacial water in the PRDX1–sulfiredoxin repair intermediate: an all-atom molecular dynamics study": SI

This file contains Supporting Figures S1-S6, Supporting Tables S1-S5, and a Supporting methods section giving every analysis algorithm in implementation-independent form. References are not included here; all citations appear in the main reference list. Items are cited at the relevant points in the main text.

#### Supporting figures

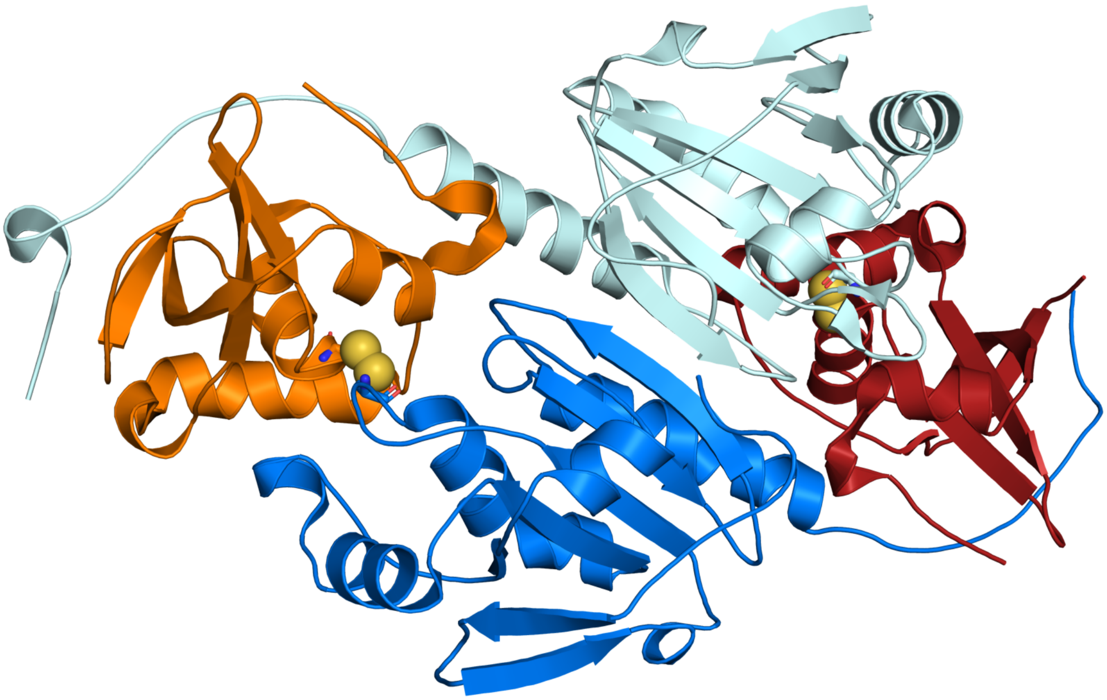

**Figure S1. The 2RII four-chain assembly.** The PRDX1 dimer (chain A, marine; chain B, pale cyan) and the two sulfiredoxin protomers (chain X, orange; chain Y, dark red), drawn as cartoons. The two PRDX1

Cys52–Srx Cys99 mixed disulfides that trap the repair intermediate are shown as sticks with their sulfur atoms as yellow spheres, one pair at each PRDX1–Srx interface.

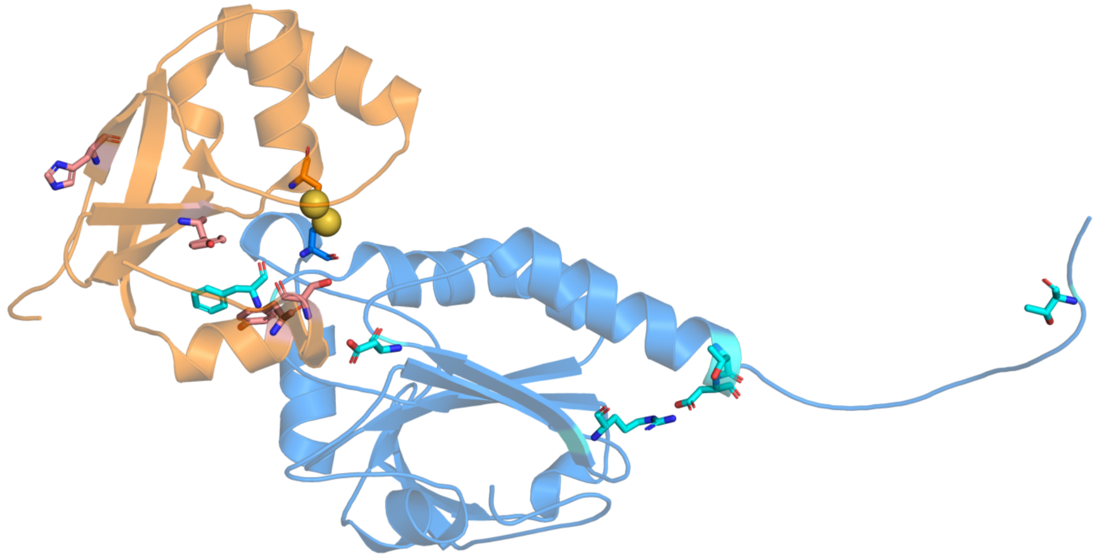

**Figure S2. Interface around the buried catalytic cysteine.** The A–X protomer pair, with PRDX1 chain A in marine and Srx chain X in orange; the cartoons are made transparent so that the side chains read clearly. Sticks show the persistent interface residues identified by the contact analysis — PRDX1 Phe50, Arg140, Asp146, Thr166, Asp167 and Thr183 with cyan carbons, and Srx His42, Tyr94, Tyr128 and Ser132 with salmon carbons. The Cys52–Cys99 mixed disulfide sits at the center of the interface, its two sulfur atoms drawn as yellow spheres; Thr183 lies out on the extended PRDX1 C-terminal arm at the right.

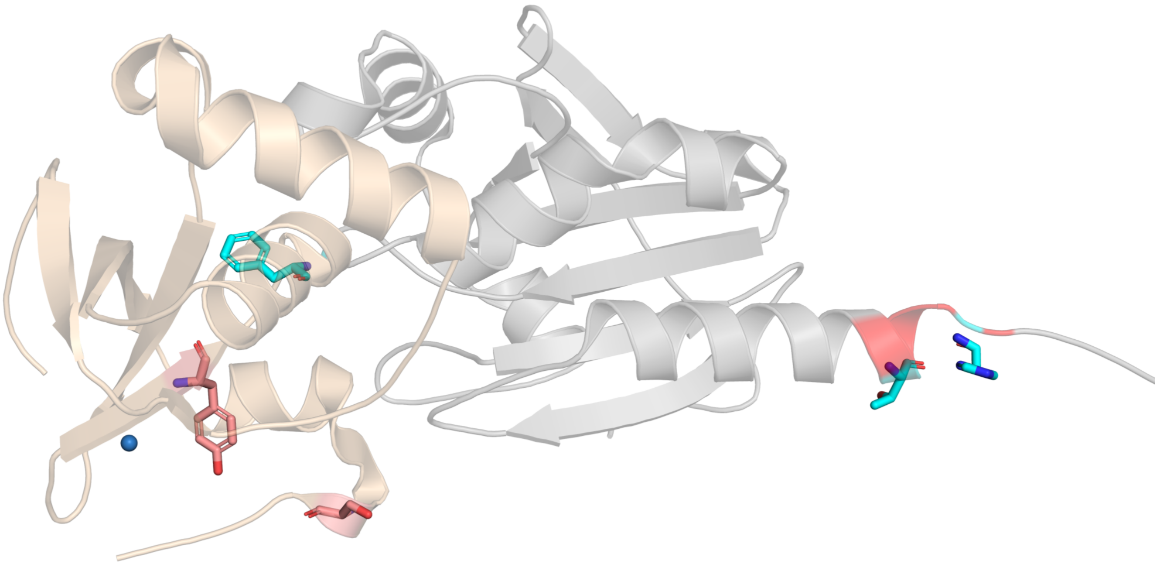

**Figure S3. The water-recognition site.** The PRDX1 165–170 region (red) at the A–X interface, with PRDX1 chain A in grey and Srx chain X in wheat. Sticks show PRDX1 Phe50, Thr166 and His169 with cyan carbons and their Srx partners Tyr94 and Ser132 with salmon carbons — the residue pairs carrying the most persistent water bridges in the simulations (Thr166–Ser132 and His169–Tyr94). Crystallographic waters within 6 Å of these residues are shown as small blue spheres.

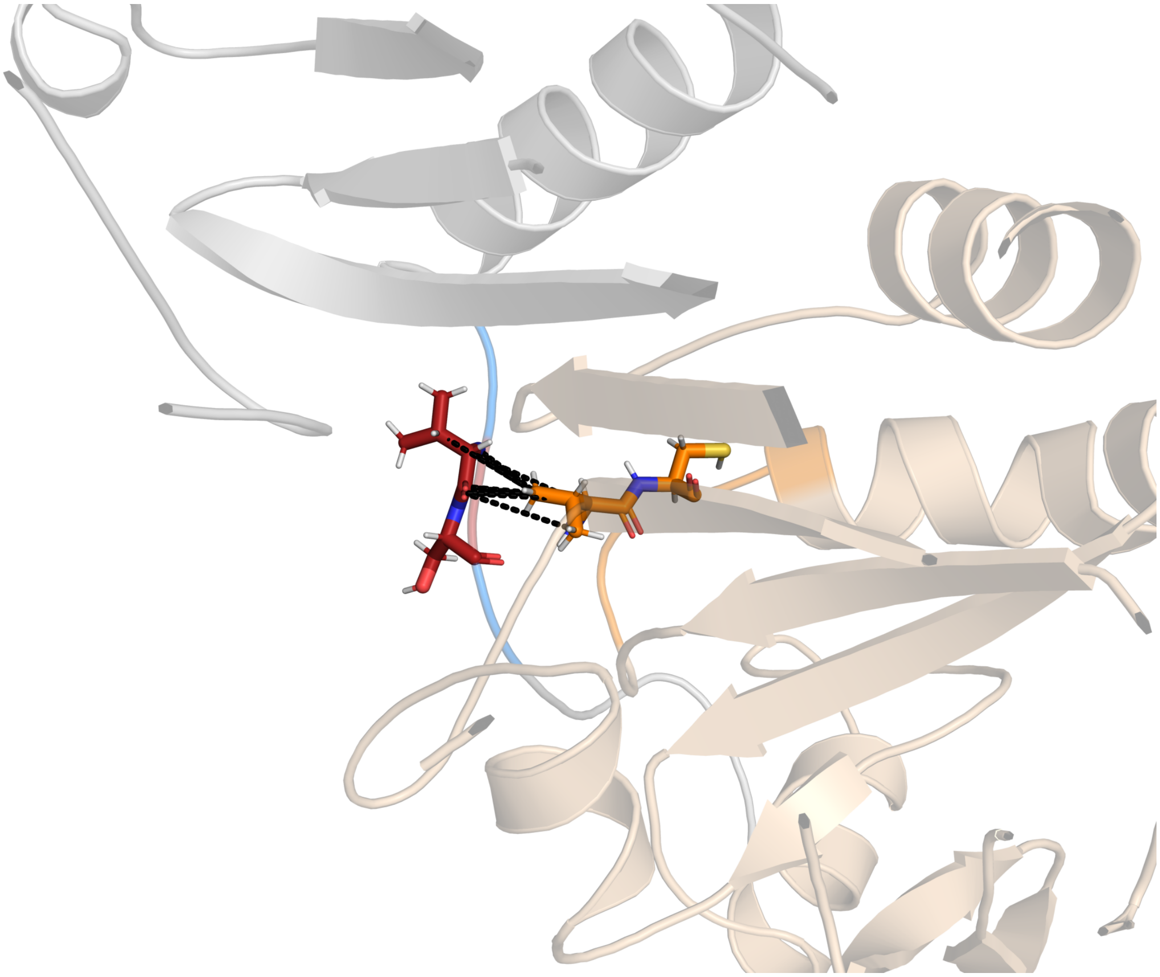

**Figure S4. The state-dependent inter-protomer contact.** A representative molecular dynamics frame of free reduced PRDX1. The C-terminal arm region of protomer A (residues 168–173, marine) packs against the peroxidatic-cysteine loop of protomer B (residues 47–53, orange), with the remainder of the two protomers in grey and wheat. Sticks show Val172 and the following residue of protomer A with firebrick carbons, and Val51 and Cys52 of protomer B with orange carbons and a yellow sulfur. Black dashes mark heavy-atom pairs within 5 Å. This is the one intra-PRDX1 pair flagged by the Srx-bound versus free comparison.

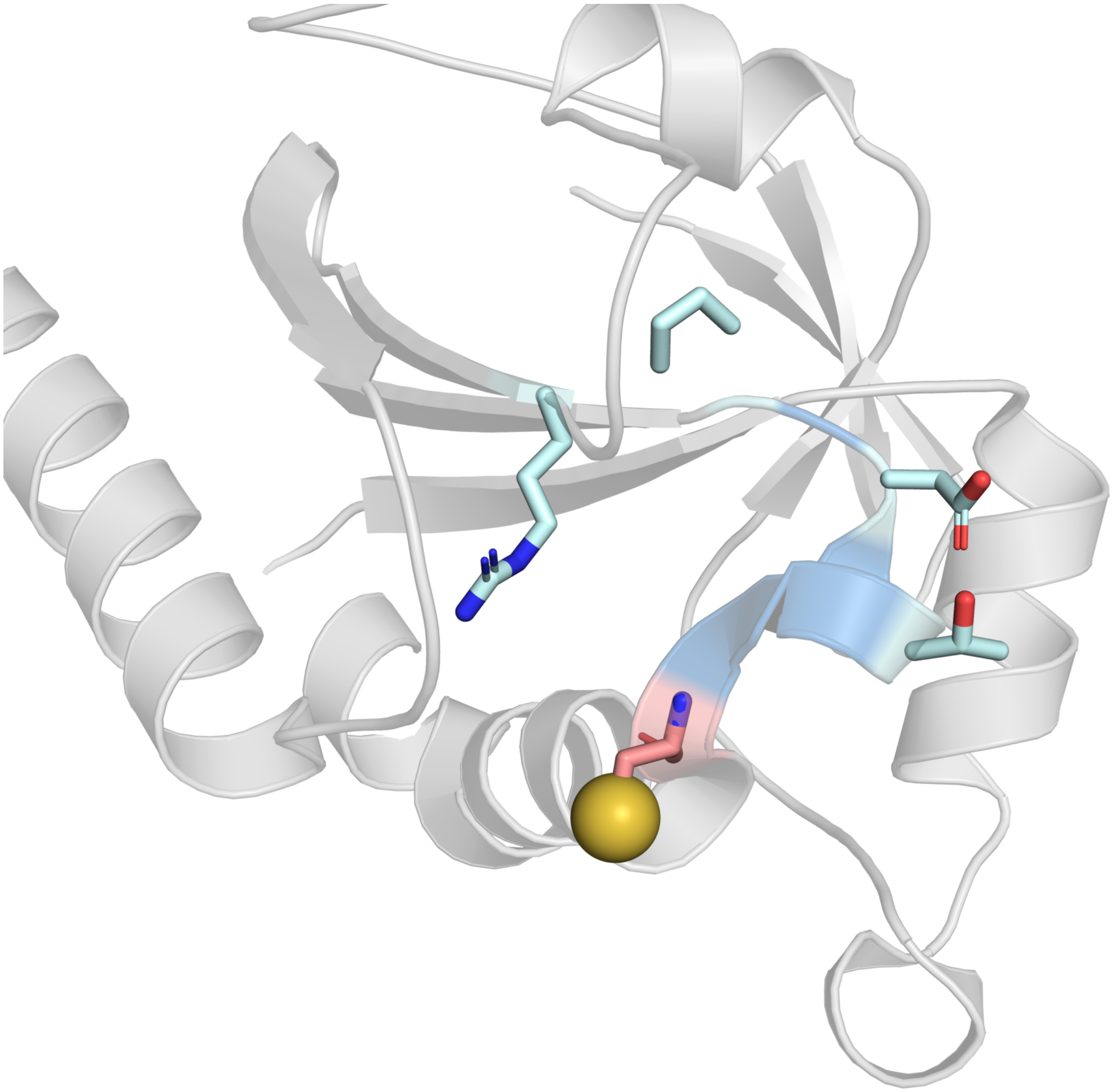

**Figure S5. The catalytic-cysteine environment.** The peroxidatic Cys52 of free PRDX1 (4XCS, chain A). The active-site loop (residues 45–52) is light blue against the grey remainder of the chain. Cys52 is drawn with salmon carbons and its SG sulfur as a yellow sphere; the surrounding Pro45, Thr47, Thr49 and Arg128 side chains have pale cyan carbons. This is the site at which the charge-state hydration analysis was carried out.

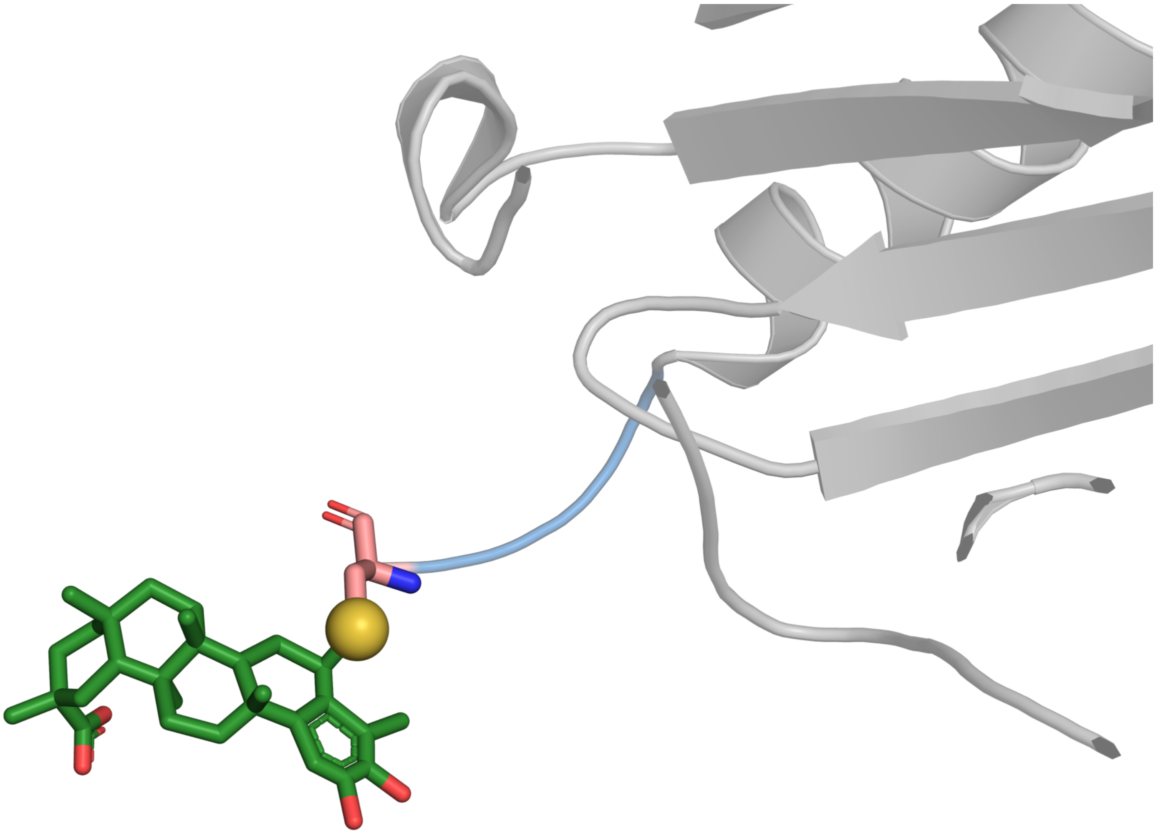

**Figure S6. The celastrol covalent adduct.** Celastrol (ligand AQR, green carbons) thioether-linked to Cys173 of PRDX1 in the 7WET-derived system. Cys173 has salmon carbons and its SG sulfur is drawn as a yellow sphere at the point of attachment; the PRDX1 C-terminal arm carrying Cys173 (residues 168–178) is light blue and the rest of the chain is grey. In this crystal structure the ligand projects away from the protein body on the flexible C-terminal arm.

### Supporting tables

**Table S1.** Cutoff-sensitivity sweep. Interface consensus contact count, number of betweenness-centrality hubs, and the hub Jaccard index versus the reference hub set, over the 27-point threshold grid (contact distance 0.40, 0.45, 0.50 nm; occupancy 0.4, 0.5, 0.6; |DCCM| 0.4, 0.5, 0.6). The reference (contact 0.45 nm, occupancy 0.5, |DCCM| 0.5) gives 149 interface contacts and hubs {ALA113, TYR116, TYR38}. Across the grid the interface consensus count ranges 83–214 and the hub Jaccard index ranges 0 to 1 (median 0.125).

| Distance (nm) | Occupancy | \|DCCM\| | Interface count | Hubs (n) | Hub Jaccard vs ref. |
|---|---|---|---|---|---|
| 0.40 | 0.4 | 0.4 | 124 | 3 | 0.000 |
| 0.40 | 0.4 | 0.5 | 124 | 5 | 0.600 |
| 0.40 | 0.4 | 0.6 | 124 | 7 | 0.111 |
| 0.40 | 0.5 | 0.4 | 103 | 2 | 0.000 |
| 0.40 | 0.5 | 0.5 | 103 | 5 | 0.333 |
| 0.40 | 0.5 | 0.6 | 103 | 6 | 0.125 |
| 0.40 | 0.6 | 0.4 | 83 | 2 | 0.000 |
| 0.40 | 0.6 | 0.5 | 83 | 6 | 0.500 |
| 0.40 | 0.6 | 0.6 | 83 | 7 | 0.111 |
| 0.45 | 0.4 | 0.4 | 172 | 1 | 0.000 |
| 0.45 | 0.4 | 0.5 | 172 | 3 | 0.500 |
| 0.45 | 0.4 | 0.6 | 172 | 6 | 0.125 |
| 0.45 | 0.5 | 0.4 | 149 | 3 | 0.000 |
| 0.45 | 0.5 | 0.5 | 149 | 3 | 1.000 |
| 0.45 | 0.5 | 0.6 | 149 | 7 | 0.111 |
| 0.45 | 0.6 | 0.4 | 130 | 3 | 0.000 |
| 0.45 | 0.6 | 0.5 | 130 | 3 | 1.000 |
| 0.45 | 0.6 | 0.6 | 130 | 6 | 0.125 |
| 0.50 | 0.4 | 0.4 | 214 | 1 | 0.000 |
| 0.50 | 0.4 | 0.5 | 214 | 3 | 0.500 |
| 0.50 | 0.4 | 0.6 | 214 | 7 | 0.111 |
| 0.50 | 0.5 | 0.4 | 188 | 1 | 0.000 |
| 0.50 | 0.5 | 0.5 | 188 | 4 | 0.400 |
| 0.50 | 0.5 | 0.6 | 188 | 5 | 0.143 |
| 0.50 | 0.6 | 0.4 | 169 | 1 | 0.000 |
| 0.50 | 0.6 | 0.5 | 169 | 4 | 0.400 |
| 0.50 | 0.6 | 0.6 | 169 | 5 | 0.143 |

*Interface counts are computed with the threshold-sweep pipeline. Across the grid the interface count ranges 91-209 and the hub Jaccard ranges 0 to 1 (median 0.5): the interface count scales monotonically with the thresholds and the occupancy-1.0 contacts of main-text Table 1 are present throughout, whereas the betweenness-hub identity is threshold-sensitive.*

**Table S2.** Per-replica convergence diagnostics for the PRDX1-Srx complex (each replica 200 ns, frames every 20 ps). Backbone RMSD is reported as the last-half mean and the final value; per-chain Calpha RMSD is the last-half mean for PRDX1 chains A and B and Srx chains C and D; Rg is the last-half radius of gyration; SG-SG lists the two mixed-disulfide distances. The two disulfides remained intact in every frame of all replicas.

| Replica | t_eq (ns) | Backbone RMSD last-half / final (Å) | Per-chain Cα RMSD A / B / C / D (Å) | Rg (Å) | SG–SG A52–C99 / B52–D99 (Å) |
|---|---|---|---|---|---|
| rep01 | 1.2 | 2.53 / 2.50 | 1.56 / 1.34 / 2.28 / 2.33 | 27.42 | 2.039 / 2.037 |
| rep02 | 1.2 | 2.72 / 2.80 | 1.98 / 1.60 / 2.79 / 2.56 | 26.78 | 2.040 / 2.038 |
| rep03 | 3.3 | 3.18 / 3.29 | 2.64 / 1.57 / 2.49 / 2.07 | 27.53 | 2.035 / 2.038 |

*Backbone RMSD and radius of gyration plateau by <=3.3 ns in all replicas (a 20 ns production discard is therefore conservative). For the slow global observables the effective sample size is N_eff = 3.6-7.3 (backbone RMSD) and 3.1-26.8 (radius of gyration), with block standard error 0.17-0.22 Å (backbone RMSD) and 0.04-0.17 Å (radius of gyration); accordingly, across-replica standard deviation (n = 3) is used as the error unit throughout the main text.*

**Table S3.** Per-system run manifests. Parameters recorded for each simulated system (values shown for the seed-1 replica; three replicas per system were run with integrator and barostat seeds 1, 2, and 3). The four systems built in OpenMM were solvated with 1.2 nm padding; the two prmtop systems were built in tleap with an isometric (iso) box. Per-replica potential energies are recorded in the deposited manifest files.

| Parameter | Complex (2RII) | Free reduced | Free thiolate | Celastrol adduct (7WET) | Cys52-sulfinate |
|---|---|---|---|---|---|
| Chains | ABXY | AB | AB | A, B (+2 AQR) | ABXY |
| Catalytic-Cys / adduct state | Cys52–Cys99 SS (×2) | Cys52 reduced | Cys52 thiolate (CYM) | Cys173–AQR thioether (×2) | Cys52 sulfinate (CSD, ×2); Srx Cys99 free thiol |
| OpenMM (platform, precision) | 8.5.1.dev-f7fa0c2 (CUDA, mixed) | 8.5.1.dev-f7fa0c2 (CUDA, mixed) | 8.5.1.dev-f7fa0c2 (CUDA, mixed) | 8.5.1.dev-f7fa0c2 (CUDA, mixed) | 8.5.1.dev-f7fa0c2 (CUDA, mixed) |
| Force field / water | amber14-all + amber14/tip3p | amber14-all + amber14/tip3p | amber14-all + amber14/tip3p | ff14SB + GAFF2 + TIP3P (tleap, iso) | ff14SB + GAFF2 ($S(=O)O^-$ as s4/o) + TIP3P (tleap, iso) |
| Box padding (nm) / mean edge (Å) | 1.2 / 112.6 | 1.2 / 111.9 | 1.2 / 111.9 | iso / 106.5 | iso / 122.8 |
| Ions $Na^+$ / $Cl^-$ | 124 / 120 | 121 / 120 | 123 / 120 | 4 / 0 | 6 / 0 |
| Waters | 43,993 | 43,973 | 43,977 | 37,777 | 57,803 |
| Nonbonded | PME, 1.0 nm, tol $5\times10^{-4}$ | PME, 1.0 nm, tol $5\times10^{-4}$ | PME, 1.0 nm, tol $5\times10^{-4}$ | PME, 1.0 nm | PME, 1.0 nm |
| Timestep (fs) / T (K) | 2 / 310.15 | 2 / 310.15 | 2 / 310.15 | 2 / 310.15 | 2 / 310.15 |
| Thermostat / barostat | Langevin 1 $ps^{-1}$ / MC 1 bar 25-step | Langevin 1 $ps^{-1}$ / MC 1 bar 25-step | Langevin 1 $ps^{-1}$ / MC 1 bar 25-step | Langevin 1 $ps^{-1}$ / MC 1 bar 25-step | Langevin 1 $ps^{-1}$ / MC 1 bar 25-step |
| Disulfide / covalent (intact) | 2 / 2 | 0 / 0 | 0 / 0 | covalent 2 / 2 | 0 / 0 (tether removed) |
| HIP residues | A84, B84 | A84, B84 | A84, B84 | A84, B84 (per prmtop) | A84, B84 (per prmtop) |
| Restrained heavy atoms | 4467 | 2880 | 2880 | 2682 | 4471 |
| Restraint schedule heat / NPT / release / free (ns) | 0.3 / 2 / 2 / 5 | 0.3 / 2 / 2 / 5 | 0.3 / 2 / 2 / 5 | 0.3 / 2 / 2 / 5 | 0.3 / 2 / 2 / 5 |
| Production (ns) / frame (ps) | 200 / 20 | 200 / 20 | 200 / 20 | 200 / 20 | 200 / 20 |
| Seeds (across replicas) | 1, 2, 3 | 1, 2, 3 | 1, 2, 3 | 1, 2, 3 | 1, 2, 3 |
| PE minimized / equilibrated ($10^6$ kJ $mol^{-1}$) | -2.352 / -1.892 | -2.327 / -1.869 | -2.331 / -1.869 | -1.905 / -1.524 | -2.934 / -2.338 |

*The celastrol-Cys173 junction (covalent SG-C6 bond, two ligands) uses approximate proxy bond, angle, and dihedral terms and is analyzed for bond persistence and ligand mobility only, not for energetic quantities. Python 3.11.15 throughout.*

**Table S4. Minimum-image separation.** For 400 evenly spaced frames per replica (every 500 ps of the 200 ns production run), molecules were made whole and the solute imaged with the largest protein molecule as anchor; the shortest distance between any solute atom and any periodic image of the solute was then evaluated over all 26 neighboring lattice translations. The relevant comparison is the 10 Å (1.0 nm) real-space cutoff used for the non-bonded interactions. Box edge and solute extent are the run-averaged and maximum values over the same frames. All three Cys52-sulfinate replicas are shown.

| System | Replica | Mean box edge (Å) | Max solute extent (Å) | Minimum solute–image distance (Å) | Frames < 10 Å (%) |
|---|---|---|---|---|---|
| PRDX1–Srx complex (2RII) | 1 | 112.6 | 110.9 | 7.5 (mean 26.6) | 1.3 |
| | 2 | 112.6 | 110.2 | 23.3 (mean 37.6) | 0 |
| | 3 | 112.5 | 107.6 | 12.9 (mean 31.5) | 0 |
| PRDX1 free, reduced | 1 | 111.9 | 83.8 | 41.2 (mean 47.4) | 0 |
| | 2 | 111.9 | 87.5 | 35.0 (mean 46.4) | 0 |
| | 3 | 111.9 | 98.6 | 24.0 (mean 44.8) | 0 |
| PRDX1 free, thiolate | 1 | 111.9 | 90.0 | 24.8 (mean 45.9) | 0 |
| | 2 | 111.9 | 98.3 | 35.1 (mean 48.5) | 0 |
| | 3 | 111.9 | 104.2 | 30.7 (mean 45.5) | 0 |
| Celastrol–Cys173 adduct (7WET) | 1 | 106.5 | 73.7 | 36.7 (mean 45.9) | 0 |
| Cys52-sulfinate complex | 1 | 122.8 | 112.9 | 11.9 (mean 34.5) | 0 |
| | 2 | 122.8 | 120.8 | 18.8 (mean 42.1) | 0 |
| | 3 | 122.8 | 116.0 | 22.8 (mean 41.0) | 0 |

Four of the five systems stay at or above 22.8 Å from their nearest periodic image in every frame examined, so none of their frames reaches the cutoff. The disulfide-linked complex is the tightest case, and the reason is geometric: its solute extent grows from 98.4 Å in the starting structure to 108–111 Å as the chain termini extend, while the box edge stays near 112.6 Å. In replica 1 the closest approach reaches 7.5 Å and 1.3% of the sampled frames fall below the cutoff; replicas 2 and 3 remain above it throughout, giving 0.4% across the three replicas. Those frames are reported here rather than discarded. The interface observables are local — contacts at 4.5 Å, water bridges at 3.5 Å and salt bridges at 4.0 Å — and are computed from the imaged coordinates, so they are evaluated well inside the separation that remains even in the closest frames. The isometric box used for the Cys52-sulfinate system carries more headroom across all three of its replicas (minima 11.9, 18.8 and 22.8 Å, none of them near the cutoff) than the cubic padding used for the four original systems.

**Table S5. Consensus interface hydrogen-bond network, disulfide-trapped versus Cys52-sulfinate.** Consensus persistent water bridges and salt bridges (occupancy ≥ 0.5, present in at least two of three replicas) were compared between the two states after normalizing the labels the two builds use: Srx is chain C/D in the OpenMM-built complex and X/Y in the tleap-built sulfinate, and Amber writes HID/HIE/HIP and CYM where the complex writes HIS and CYS. Because the assembly is a homodimer with two equivalent interfaces, pairs are counted protomer-agnostically, so the 20 and 36 chain-specific bridges of the two states reduce to 14 and 21 distinct residue pairs.

| residue pairs | water bridges | salt bridges |
|---|---|---|
| present in both states | 11 | 1 |
| only in the disulfide-trapped complex | 3 | 1 |
| only in the Cys52-sulfinate complex | 10 | 0 |
| Jaccard index | 0.46 | 0.50 |

Shared water-bridge pairs: His169–Tyr94, Thr166–Ser132, Trp177–Asp91, Leu147–Tyr128, Glu123–Arg126, Glu123–Ser123, Lys120–Arg126, Ala175–Asp91, Gly176–Asp91, His169–Arg51 and Val172–Tyr92. Present only with the disulfide: Asp146–Ala131, Lys185–Asp74 and Thr166–Pro52. Present only with the sulfinate: Cys52–Cys99, Cys52–His100, Cys52–Phe96, Cys52–Gly98, Phe50–Asp80, Phe50–Gly97, Phe50–Phe96, Arg140–Ser132, Asp146–Ser132 and Gly170–Ser132 — the first four of these occupy the position vacated by the engineered bond. The single shared salt bridge is Glu123–Arg126; Lys185–Asp74 is present only with the disulfide.

#### Supporting methods: analysis algorithms

Every analysis reported in this work is specified below in implementation-independent form, with all thresholds and controls given explicitly, so that the results can be reproduced from the description alone. Distances are heavy-atom distances unless stated otherwise. “Occupancy” always means the fraction of analyzed frames in which a feature is present. Analyzed frames are the post-equilibration frames of each replica: the first 20 ns of each 200 ns production run is discarded and every fifth remaining frame is used (40 ps spacing). “Consensus” always means present in at least two of the three replicas, and is reported as the mean occupancy ± the across-replica standard deviation.

##### A1. Persistent interface and intra-protein contacts

```
for each replica r:
   P <- residues of PRDX1 chains, S <- residues of Srx chains        # by author chain, not
   topology index
   candidates <- residue pairs whose mean C-alpha separation < 1.6 nm
   for each frame f, for each candidate pair (i,j):
      contact(i,j,f) <- [ min heavy-atom distance(i,j,f) < 0.45 nm ]
   occupancy_r(i,j) <- mean over frames of contact(i,j,.)
   persistent_r <- { (i,j) : occupancy_r(i,j) >= 0.5 }
   interface_r  <- { (i,j) in persistent_r : i in P xor j in P }
   intra_r      <- { (i,j) in persistent_r : i in P and j in P }
consensus <- pairs appearing in >= 2 of the 3 replica sets
report mean occupancy +/- SD across the replicas in which the pair appears
```

##### A2. Interface water bridges

```
for each replica r, for each frame f:
   for each water W:
      A <- polar atoms (N,O) of PRDX1 within 0.35 nm of W.O
      B <- polar atoms (N,O) of Srx   within 0.35 nm of W.O
      if A empty or B empty: continue                     # must contact both sides
      if not exists X in (A union B) such that W donates a hydrogen bond to X
           with angle(O-H...X) >= 120 deg and distance(H...X) <= 0.25 nm: continue
      record the bridge by RESIDUE-PAIR identity (residue of A, residue of B),
         not by the identity of the water, so exchange of the bridging water is allowed
   occupancy_r(pair) <- fraction of frames in which the pair is bridged by any water
persistent_r <- { pair : occupancy_r(pair) >= 0.5 };  consensus as in A1
```

##### A3. Salt bridges

```
charged donors    <- side-chain N of Arg, Lys, His(+)
charged acceptors <- side-chain O of Asp, Glu
salt bridge in frame f  <-  any donor-acceptor distance <= 0.40 nm
collapse atom pairs to residue pairs before computing occupancy
persistent and consensus as in A1
```

##### A4. Correlated motion and network hubs

```
align each chain onto its own frame 0 separately          # removes inter-chain tumbling
d(i,f) <- displacement of C-alpha of residue i in frame f from its mean position
DCCM(i,j) <- <d(i,.).d(j,.)> / sqrt(<|d(i,.)|^2><|d(j,.)|^2>)     # intra-PRDX1 only
reproducibility: Pearson r on the upper triangle, split-half within a replica
                          and between replicas
graph G: nodes = PRDX1 residues
              edge (i,j) iff occupancy(i,j) >= 0.5 AND |DCCM(i,j)| > 0.5    # no fallback rule
hubs <- top 15 residues by betweenness centrality of G, pooled over the homodimer
```

##### A5. Srx-bound versus free comparison, with two null models

```
equalize: draw the same number of frames (1500) per state per replica
for each protomer-agnostic residue pair (i,j):
   delta(i,j) <- occupancy_bound(i,j) - occupancy_free(i,j)
permutation null:  shuffle the bound/free labels across replicas 3000 times,
                         recompute delta, take the 95th percentile of |delta| -> T_perm
replicate floor:   |delta| between two replicas of the SAME state,
```

```
                              95th percentile -> T_rep        # what noise alone produces at n =
                              3
report (i,j) as differing only if:
        |delta(i,j)| > T_perm  AND  |delta(i,j)| > T_rep
        AND occupancy >= 0.5 in at least one of the two states
```

##### A6. Charge-state hydration and the matched-count control

```
for each Cys52 SG, per frame: count waters with O within 3.5 / 4.5 / 6.0 / 8.0 A
dipole orientation <- cos(angle between the water dipole and the SG->O vector)
                                   negative => hydrogens point toward the sulfur
clustering: build the first-shell water graph, edge iff O-O < 0.35 nm
                   fLC <- (size of largest connected component) / (number of first-shell waters)
MATCHED-COUNT CONTROL (this is the control that matters):
   bin frames by first-shell water count k
   for each k present in BOTH states: delta_fLC(k) <- fLC_thiolate(k) - fLC_thiol(k)
   report the count-weighted mean of delta_fLC(k)
   -> a difference that survives is a change in network organization;
        one that vanishes was only a difference in how many waters were present
```

##### A7. Minimum-image separation

```
make molecules whole, then image the solute with the LARGEST protein molecule as anchor
   (without an anchor the separately-bonded halves drift into different periodic images)
for each analyzed frame:
   for each of the 26 neighboring lattice translations t of the box vectors:
      d(t) <- min over solute atom pairs (a,b) of | pos(a) - (pos(b) + t) |
   min_image(frame) <- min over t of d(t)
report the per-replica minimum and mean, and the fraction of frames with
min_image < the 1.0 nm real-space cutoff
```
